# Structural insights into human propionyl-CoA carboxylase (PCC) and 3-methylcrotonyl-CoA carboxylase (MCC)

**DOI:** 10.1101/2024.04.30.591959

**Authors:** Fayang Zhou, Yuanyuan Zhang, Yuyao Zhu, Qiang Zhou, Yigong Shi, Qi Hu

## Abstract

Propionyl-CoA carboxylase (PCC) and 3-methylcrotonyl-CoA carboxylase (MCC) are biotin-dependent carboxylases (BDCs) that catalyze the metabolism of odd-chain fatty acids, cholesterol, and specific amino acids. For human PCC and MCC, only a low-resolution (15 Å) three-dimensional structure of human PCC has been reported. Here, we report high-resolution (2.29–3.38 Å) cryo-EM structures of human PCC and MCC holoenzymes in their apo and acetyl-CoA and propionyl-CoA-bound states. Propionyl-CoA and acetyl-CoA bind to PCC with almost identical binding modes, indicating that the acyl-CoA specificity of PCC is largely attributed to minor differences in interactions mediated by the acyl groups. In MCC, biotin is relocated from an exo-site to an endo-site upon acetyl-CoA binding, suggesting coordination between biotin binding and acyl-CoA binding. Our work provides insights into the substrate specificity and catalytic process of BDCs.

## Introduction

Biotin-dependent carboxylases (BDCs) catalyze the carboxylation of several key metabolites using covalently linked biotin as a cofactor(*1, 2*). In humans, there are four types of BDCs: acetyl-CoA carboxylase (ACC), propionyl-CoA carboxylase (PCC), 3-methylcrotonyl-CoA carboxylase (MCC), and pyruvate carboxylase (PYC)(*1, 2*). All BDCs contain a biotin carboxylase (BC) domain, a carboxyltransferase (CT) domain, and a biotin carboxyl carrier protein (BCCP) domain(*1, 2*). Among the four human BDCs, ACC and PYC each consist of a single chain that incorporates the BC, CT, and BCCP domains(*2*). In contrast, PCC and MCC have their BC, CT, and BCCP domains divided into two chains: the α subunits contain the BC and BCCP domains, with an intermediate BC-CT interaction (BT) domain between them, while the β subunits solely consist of the CT domain(*2*).

In humans, PCC and MCC catalyze the carboxylation of propionyl-CoA and 3-methylcrotonyl-CoA (MC-CoA), respectively(*3, 4*). Propionyl-CoA is an intermediate in the metabolism of odd-chain fatty acids, cholesterol, and amino acids including valine, isoleucine, methionine, and threonine; its carboxylated product (S)-methylmalonyl-CoA is then converted into succinyl-CoA and subsequently metabolized in the tricarboxylic acid cycle(*3*). MC-CoA is an intermediate in the metabolism of leucine; following carboxylation, the resulting product 3-methylglutaconyl-CoA (MG-CoA) is further converted into 3-hydroxy-3-methylglutaryl-CoA (HMG-CoA) and then involved in the synthesis of cholesterol(*5*). Therefore, both PCC and MCC play important roles in metabolic processes. Loss-of-function mutations in MCC and PCC lead to severe metabolic disorders(*3, 6–8*).

The protein sequences and functions of PCC and MCC are conserved from bacteria to humans. Structures of bacteria PCC and MCC holoenzymes reveal how the α subunits and β subunits assemble to form the holoenzymes(*9, 10*). A crystal structure of a chimeric bacteria PCC holoenzyme, obtained by co-expressing the α subunit of *Ruegeria pomeroyi* PCC (RpPCC) with the β subunit of *Roseobacter denitrificans* PCC (RdPCC) in *Escherichia coli*, was determined at a resolution of 3.2 Å (*9*). The PCC holoenzyme is a dodecamer formed by six α subunits and six β subunits, thereby referred to as the α_6_β_6_ holoenzyme(*9*). Crystal structures of a bacteria MCC (*Pseudomonas aeruginosa* MCC, PaMCC) holoenzyme reveal that the MCC holoenzyme is also a α_6_β_6_dodecamer(*10*).

Studies of bacteria BDCs also reveal the catalytic mechanism of BDCs. The BDC catalyzed carboxylation reaction involves a two-step process(*1, 2*). In the first step, the biotin covalently linked to the BCCP domain is carboxylated in the BC domain, using bicarbonate as the donor of the carboxyl group(*11*). A crystal structure of *Escherichia coli* BC in complex with its substrates indicates that E296 (equivalent to E357 and E343 in human PCCα and MCCα, respectively) acts as a base to deprotonate bicarbonate(*12*). The deprotonated bicarbonate attacks the γ-phosphate of ATP to form carboxyphosphate, which is then decomposed into CO_2_ and a phosphate group. The phosphate group is proposed to deprotonate biotin to form an enolate biotin intermediate. Another key residue R338 (equivalent to R399 and R385 in human PCCα and MCCα, respectively) stabilizes this intermediate to facilitate its reaction with CO_2_ to form carboxylated biotin(*12*). In the second step, the carboxylated biotin is carried to the CT domain by the BCCP domain(*13*). In the CT domain, the carboxyl group is transferred from the biotin to acetyl-CoA, propionyl-CoA, MC-CoA, or pyruvate(*2, 14*). Crystal structures of the β subunit of *Streptomyces coelicolor* PCC (ScPCC) shows that the CT domains dimerize to form the acyl-CoA and biotin binding pockets(*15*). G182 and G183 in one ScPCC CT domain (equivalent to G202 and G203 in human PCCβ and A218 and G219 in human MCCβ) form an oxyanion hole for binding to the carbonyl group of the acyl moiety, while G419 and A420 from the other ScPCC CT domain (equivalent to G437 and A438 in human PCCβ and A447 and G448 in human MCCβ) also form an oxyanion hole to bind to the biotin carbonyl group. The reaction is initiated by decarboxylation of the carboxylated biotin, resulting in free CO_2_ and activated biotin. The activated biotin deprotonates the C-α of the acyl moiety, which then attacks the free CO_2_ to form carboxylated acyl-CoA(*15*).

In contrast to the extensive studies of bacterial BDCs, our understanding of human PCC and MCC is limited by the lack of high-resolution three-dimensional structures. Thus far, only a low-resolution cryo-EM structure of human PCC holoenzyme (15 Å) has been reported(*9*), highlighting the need for high-resolution structural information of human PCC and MCC.

In this study, we purified the endogenous PCC and MCC from human cells and determined the cryo-EM structures of human PCC and MCC holoenzymes in the apo (PCC or MCC holoenzyme alone), acetyl-CoA-bound, and propionyl-CoA-bound states, at resolutions ranging from 2.29 to 3.38 Å. Structural analysis reveals that human PCC and MCC holoenzymes have overall structures similar to their bacterial homologs. For human PCC holoenzyme, the structure in the acetyl-CoA-bound state closely resembles the structures in the apo and the propionyl-CoA-bound state. In these structures, the covalently linked biotin binds to a site (exo-site) that is distant from the catalytic residues in the CT domain, rendering it catalytically incompetent. For human MCC holoenzyme, the covalently linked biotin also binds to an exo-site in the apo structure. However, when MCC binds to acetyl-CoA, but not propionyl-CoA, it adopts a different conformation in its acyl-CoA binding pocket compared to the apo MCC, and the covalently linked biotin is located to an endo-site that is closer to the catalytic residues in the CT domain.

## Results

### Overall structures of human PCC and MCC holoenzymes

The BDCs endogenously expressed in Expi 293F cells were purified using streptavidin beads (the Strep-Tactin^®^XT resin) since streptavidin can specifically bind to the covalently linked biotin in the BDCs. The purified BDCs were further purified using size exclusion chromatography. Mass spectrometry analysis identified all four types of human BDCs in the purified samples (fig. S1). The samples were subjected to cryo-EM sample preparation (fig. S1, A to C). We did not characterize the enzyme activities of the mixed BDCs because the current methods used to evaluate the carboxylase activities of BDCs, such as measuring the ATP hydrolysis or incorporation of radio-labeled CO_2_, are unable to differentiate the specific carboxylase activity of each BDC.

The cryo-EM structures of the PCC and MCC holoenzymes alone, referred to as PCC-apo and MCC-apo, were determined at resolutions of 3.02 Å and 2.29 Å, respectively (figs. S2A, S3, A to D and S4, A to D, tables S1 and S2). The cryo-EM maps have good quality to support model building (figs. S5, S6 and S7).

Both the PCC and MCC holoenzymes consist of six α subunits and six β subunits. The cryo-EM density of the BC domains in one layer of three MCCα subunits in the MCC-apo structure is missing. To demonstrate the complete structure of MCC holoenzyme, these domains were built into the structure based on the structure of BC domains in the other layer of MCCα subunits (fig. S8D). The α_6_β_6_dodecamer can be divided into four layers. The six β subunits are stacked into two layers to form the core of the holoenzymes, with three β subunits laid head to end in each layer (Fig. 1, A and B). The six α subunits are separated into two layers and situated at the top and bottom sides of the β subunit core, with each layer contains three α subunits. A major difference between the PCC and MCC holoenzymes is that the three α subunits in each layer have no contact in the PCC holoenzyme but interact with each other in the MCC holoenzyme (Fig. 1, A and B).

**Fig. 1.**
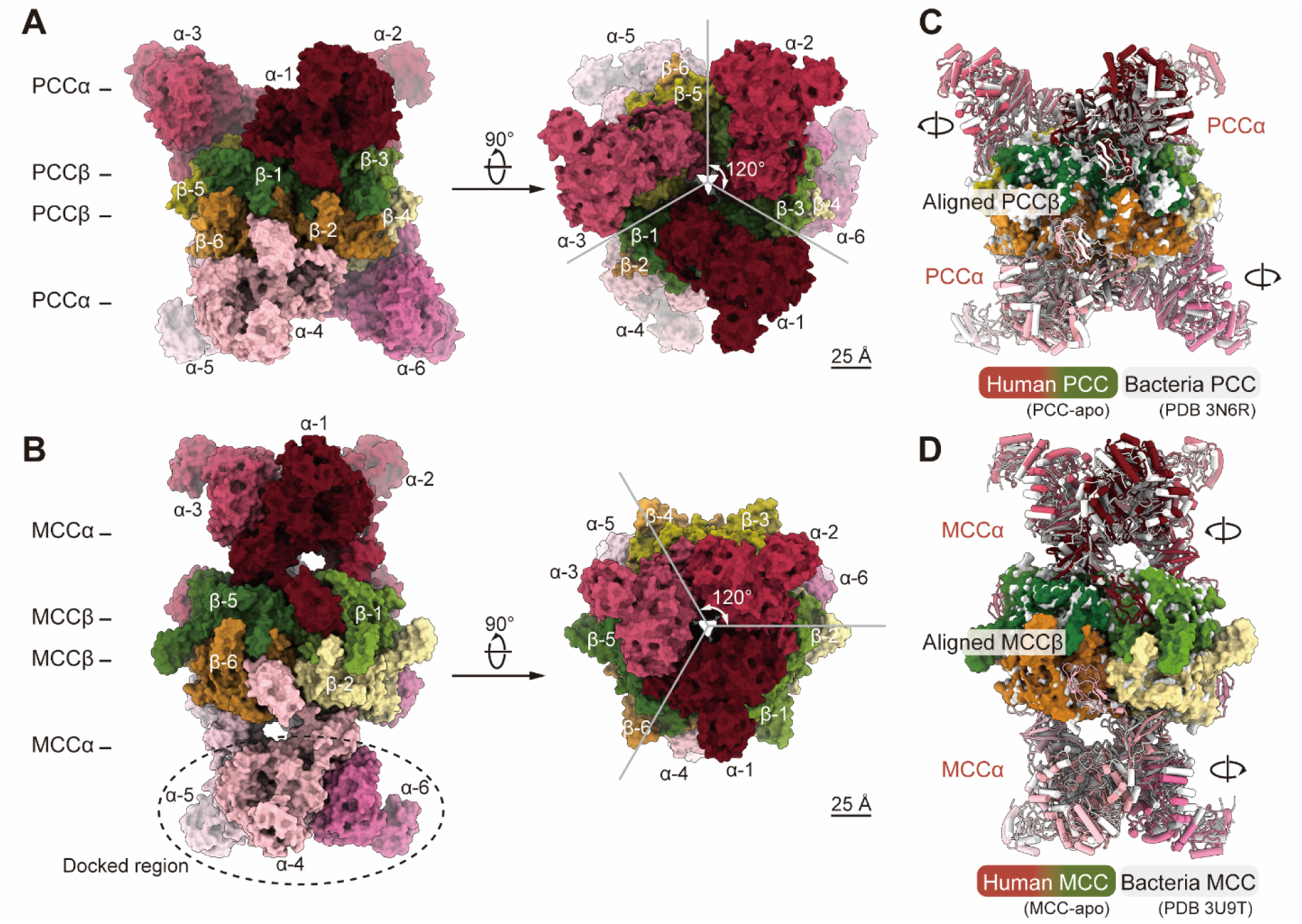
The overall structures of human PCC and MCC holoenzymes. (**A, B**) The side view (left) and top view (right) of human PCC holoenzyme (PCC-apo) (**A**) and MCC holoenzyme (MCC-apo) (**B**). Both holoenzymes are dodecamers consisting of six α subunits and six β subunits. The BC domains in one layer of MCCα subunits in the MCC-apo structure (marked by a dashed ellipse) are not resolved in the cryo-EM map, so they were docked into the structure according to the structure of BC domains in the other layer of MCCα subunits (see fig. S8). (**C**) Alignment of the PCCβ core in human PCC holoenzyme (PCC-apo) with that in a bacteria PCC holoenzyme (PDB code: 3N6R)(*9*). (**D**) Alignment of the MCCβ core in human MCC holoenzyme (MCC-apo) with that in a bacteria MCC holoenzyme (PDB code: 3U9T)(*10*). The β subunit cores in human PCC and MCC holoenzymes closely resemble their bacterial homologs, whereas the α subunits exhibit slightly different orientations in comparison to their bacterial homologs.

The overall structures of human PCC and MCC holoenzymes are similar to the reported crystal structures of bacteria PCC and MCC holoenzymes, respectively (Fig. 1, C and D). Specifically, alignment of the β subunits in the PCC-apo structure with those in the structure of a chimeric bacteria PCC holoenzyme (PDB code: 3N6R) shows that the β subunit cores in the two structures are very similar(*9*). Each α subunit in human PCC holoenzyme has a minor rotation relative to the corresponding α subunit in the bacteria PCC holoenzyme (Fig. 1C). Similarly, the β subunit core in the MCC-apo structure aligns well with that in the crystal structure of a bacteria MCC holoenzyme (PDB code: 3U9T), with each α subunit slightly rotated in comparison to the corresponding α subunit in the bacteria MCC holoenzyme(*10*) (Fig. 1D). The structures of the BC, BT, and BCCP domains in the α subunits of human PCC and MCC resemble the corresponding domains in bacteria PCC and MCC(*9, 10*), except that the BT domain of human MCC has one more β strand in comparison to that of the PaMCC (fig. S9).

### Organization of the catalytic domains in human PCC and MCC holoenzymes

We incubated the purified human BDCs with propionyl-CoA (fig. S1, D to G), which is the substrate of PCC, and determined the cryo-EM structures of the propionyl-CoA-bound PCC and MCC holoenzymes (referred to as PCC-PCO and MCC-PCO), at resolutions of 2.80 Å and 2.36 Å, respectively (figs. S2B, S3, E to H, S4, E to H, S6 and S7, tables S1 and S2). The BC domains in one layer of three MCCα subunits in the MCC-PCO structure were not resolved in the cryo-EM map, and they were docked into the structure according to the structure of BC domains in the other layer of MCCα subunits (fig. S8E). The structures of PCC-PCO and MCC-PCO are almost identical to the structures of PCC-apo and MCC-apo, respectively. Although bicarbonate, MgCl_2_, and ATP were added to the purified BDC samples to facilitate the carboxylation reaction, the cryo-EM density of the biotin region does not provide conclusive evidence as to whether the covalently linked biotin in the PCC holoenzyme structures was indeed carboxylated. Thus, the cryo-EM structures only contain the uncarboxylated biotin.

The two-step carboxylation reactions catalyzed by PCC and MCC require the coordination of the BC, BCCP domains in the α subunit and the CT domain in the β subunit (Fig. 2A). In human PCC holoenzyme, each α subunit binds to one β subunit containing the CT domain that can be divided into CT-N and CT-C subdomains (Fig. 2B). The BC domain in the α subunit is situated on the CT-C subdomain, while the BCCP domain also attaches to the CT-C subdomain and positions the covalently linked biotin to a pocket adjacent to the propionyl-CoA binding pocket. The BT domain tightly interacts with both the CT-N and CT-C subdomains and functions as a hub to hold the BC and BCCP domains together with the CT domain (Fig. 2B).

**Fig. 2.**
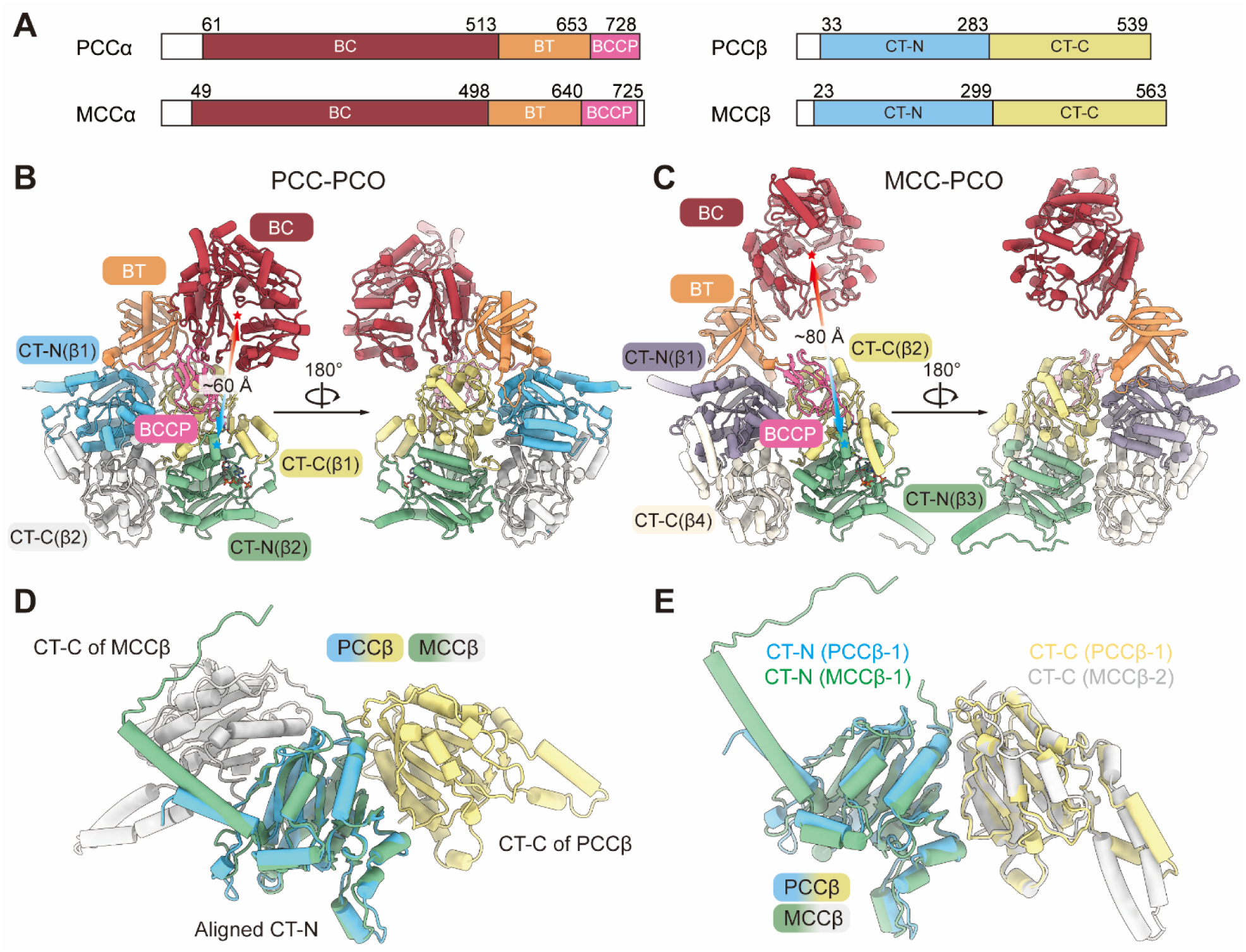
Domain organization of human PCC and MCC holoenzymes. (**A**) Schematic illustration of the domain organization of human PCC (top) and MCC (bottom). Both PCC and MCC consist of α and β subunits. From the N-terminal to the C-terminal, each α subunit contains a BC (red), a BT (orange), and a BCCP (pink) domain. Each β subunit contains a CT domain, which can be divided into the CT-N (blue) and CT-C (yellow) subdomains. (**B, C**) Organization of the catalytic domains in the propionyl-CoA-bound PCC holoenzyme (PCC-PCO) (**B**) and MCC holoenzyme (**C**). (**D**) Alignment of the CT-N subdomain of PCCβ with that of MCCβ shows the different arrangement of the CT domains in human PCC and MCC. (**E**) One PCCβ subunit aligns well with the CT-N and CT-C subdomains from two adjacent MCCβ subunits in the MCC holoenzyme.

In contrast, in human MCC holoenzyme, each α subunit binds to two β subunits (referred to as β1 and β2) (Fig. 2C). The CT-N subdomain of β1 interacts with the CT-C subdomain of β2. The BT domain of the α subunit associates with both the CT-N and CT-C subdomains of β1, while the BCCP domain attaches to the CT-C subdomain of β2. The propionyl-CoA binds to a pocket formed by the CT-C of β2 and the CT-N of a third β subunit (β3) from the other β subunit layer. The BC domain has no contact with the β subunits.

The variances in the organization of the catalytic domains in the PCC holoenzyme and the MCC holoenzyme are partly attributed to the distinct arrangement of their CT domains (Fig. 2D). Alignment of the CT-N subdomains shows that the CT-C subdomains in PCC and MCC have distinct orientations (Fig. 2D). The CT domain in PCC can perfectly align with a combination including the CT-N of β1 and the CT-C of β2 of MCC (Fig. 2E).

### Propionyl-CoA binding in human PCC holoenzyme

In each PCC holoenzyme, six propionyl-CoA molecules are observed binding to the six active sites of CT domains. Each binding site for propionyl-CoA is formed by the CT-C subdomain of one β subunit and the CT-N subdomain of another β subunit in the other layer (Fig. 3A). The conformation of the propionyl-CoA binding pocket in the PCC-PCO structure is only slightly different from that in the PCC-apo structure (Fig. 3B). The AMKM motif in the BCCP domain and the biotin covalently linked to it also show only minor conformational differences in the PCC-PCO structure compared to the PCC-apo structure (Fig. 3C). The distances between the ureido ring carbonyl oxygen of the biotin and the main chain amide of the catalytic residues G437 and A438 in the corresponding CT domain in the PCC-PCO structure are more than 7 Å, which is too far for an efficient catalytic process (Fig. 3C). This finding suggests that the biotin may need to relocate to a site that is closer to the catalytic residues to facilitate the carboxylation reaction.

**Fig. 3.**
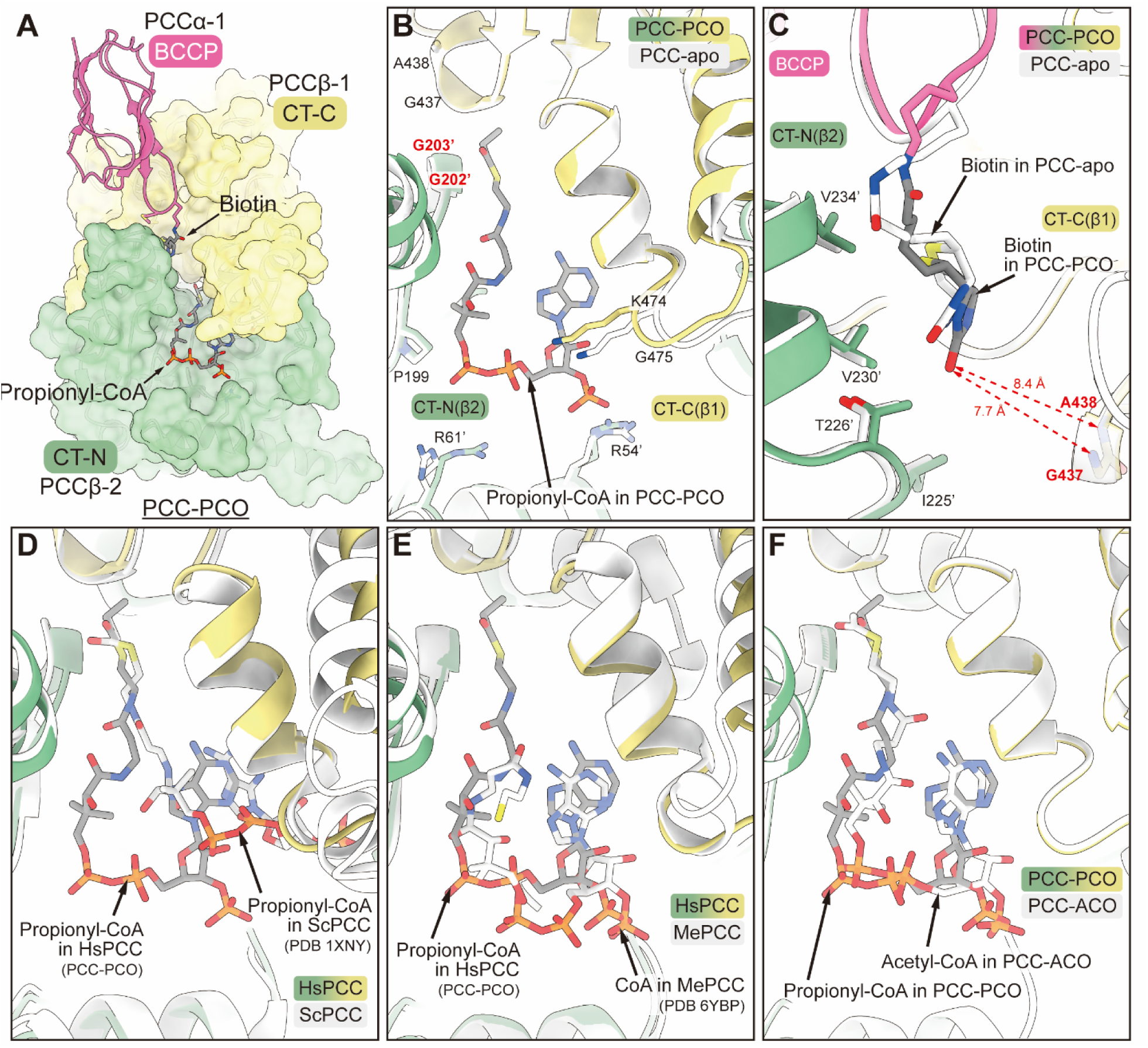
Acyl-CoA binding in human PCC holoenzyme. (**A**) Surface representation of the binding pockets of biotin and propionyl-CoA in the PCC-PCO structure. (**B, C**) Alignment of the CT-N subdomain in the PCC-PCO structure with that in the PCC-apo structure. The conformation of the acyl-CoA binding pocket in the PCC-PCO structure is similar to that in the PCC-apo structure (**B**), and the AMKM motif and biotin covalently linked to it show minor conformational differences (**C**). The distances between the carbonyl oxygen in the ureido ring of the biotin and the main chain amide of the catalytic residues G437 and A438 in the PCC-PCO structure are 7.7 Å and 8.4 Å, respectively. (**D**) Alignment of the CT-N subdomain in the PCC-PCO structure with that in the ScPCC structure (PDB code: 1XNY)(*15*). The propionyl-CoA in the two structures has distinct binding modes. (**E**) Alignment of the CT-N subunit in the PCC-PCO structure with that in the MePCC structure (PDB code: 6YBP)(*16*). The propionyl-CoA binds at the similar positions in both structures. (**F**) Alignment of the CT-N subunit in the PCC-PCO structure with that in the PCC-ACO structure. The propionyl-CoA and acetyl-CoA in the two structures show highly similar binding modes.

Before our study, two structures were reported to demonstrate the binding of CoA or propionyl-CoA in the CT domains of bacteria PCCs. One is the crystal structure of the β subunits of ScPCC in complex with propionyl-CoA (PDB code: 1XNY)(*15*). The other is the cryo-EM structure of the *Methylorubrum extorquens* PCC (MePCC) holoenzyme in complex with CoA (PDB code: 6YBP)(*16*). The binding mode of propionyl-CoA in our PCC-PCO structure is distinct from that in the ScPCC β subunit structure, but similar to that in the MePCC holoenzyme structure (Fig. 3D, E).

### Acetyl-CoA binding in human PCC and MCC holoenzymes

In another study from our group(*17*), we incubated the purified human BDCs with acetyl-CoA, which is a substrate of ACC, and determined the cryo-EM structure of the acetyl-CoA-bound ACC. Using the same batch of purified BDCs (fig. S1, H to K), we also determined the cryo-EM structures of human PCC and MCC holoenzymes in the acetyl-CoA-bound state (referred to as PCC-ACO and MCC-ACO) at resolutions of 3.38 Å and 2.85 Å, respectively (figs. S2C, S3, I to L, S4, I to L, S6 and S7, tables S1 and S2). The BC domains of one PCCα subunit in the PCC-ACO structure and three MCCα subunits in the MCC-ACO structure are not resolved. These domains were built into the structures based on the resolved BC domains in other α subunits (fig. S8, C and F).

The PCC-ACO structure closely resembles the PCC-apo and the PCC-PCO structures. The acetyl-CoA in the PCC-ACO structure has a binding mode similar to the propionyl-CoA in the PCC-PCO structure (Fig. 3F). The conformation of the acyl-CoA binding pocket in both structures is also highly similar (Fig. 3F). It was reported that D422 in the β subunit of a bacteria PCC (ScPCC) is crucial for the substrate specificity of ScPCC, and the D422I mutation significantly increased the catalytic activity of ScPCC towards acetyl-CoA(*15, 18*). But the PCC-PCO structure shows that D440, which corresponds to D422 in the β subunit of ScPCC, is not directly involved in the interaction with propionyl-CoA or acetyl-CoA (Fig. 3F).

The overall structure of MCC-ACO is also similar to that of MCC-apo, but alignment of the CT-N subdomains shows that the position of the BCCP domain and the conformation of the acyl-CoA binding pocket in the MCC-ACO structure are significantly different from those in the MCC-apo structure (Fig. 4A). Specifically, the BCCP domain moves towards the CT-C subdomain in the MCC-ACO structure compared to that in the MCC-apo structure (Fig. 4A). Consequently, the covalently linked biotin moves from an exo-site to an endo-site that is deeper in the pocket formed by the CT-N and CT-C subdomains from two β subunits (Fig. 4B). At the exo-site, the ureido ring carbonyl oxygen of the biotin is about 8 Å away from the main chain amide of the catalytic residues G437 and A438 (fig. S10A). In contrast, at the endo-site, the distance is shortened to about 3.5 Å (fig. S10B). The acyl-CoA binding pocket, also formed by the CT-N and CT-C subdomains from two β subunits, switches from an open conformation in the MCC-apo structure to a closed conformation in the MCC-ACO structure (Fig. 4, C to E), with the major conformational change occurring at two helices (residues 474 to 517) in the CT-C subdomain and one helix (residues 243 to 252) in the CT-N subdomain (Fig. 4C).

**Fig. 4.**
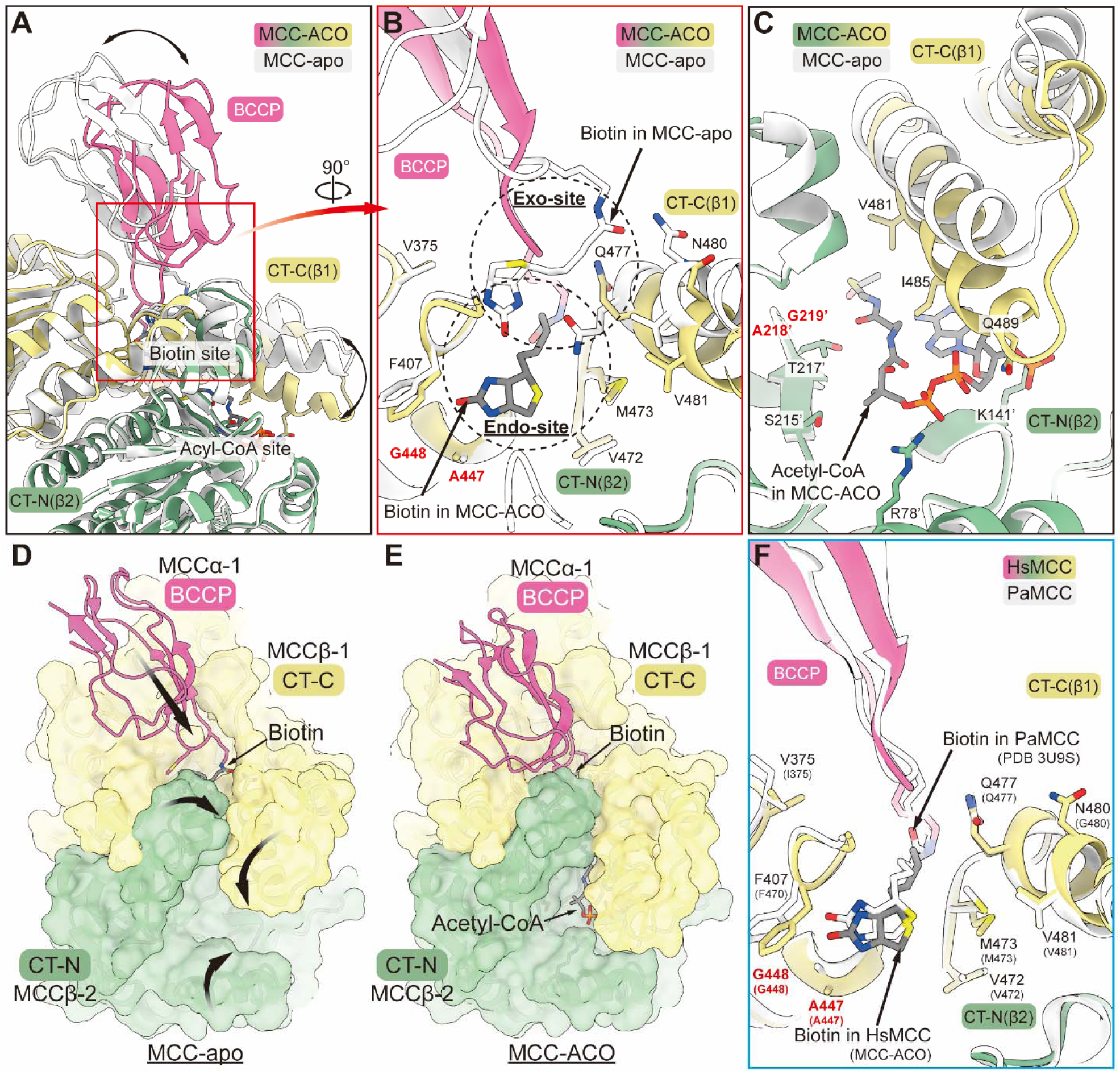
Conformations of human MCC holoenzyme in apo and acyl-CoA-bound states. (**A**– **C**) Alignment of the CT-N subdomain in the MCC-ACO structure with that in the MCC-apo structure. The BCCP domain moves towards the CT-C subdomain in the MCC-ACO structure compared to that in the MCC-apo structure (**A**). The covalently linked biotin binds to an exo-site in the MCC-apo structure but relocates to an endo-site that is closer to the catalytic residues A447 and G448 (see fig. S10) in the MCC-ACO structure (**B**). The helices around the acyl-CoA binding pocket show conformational differences in the MCC-ACO structure compared to that in the MCC-apo structure (**C**). (**D**, **E**) Surface representation of the binding pockets of biotin and acyl-CoA in the structures of MCC-apo and MCC-ACO. The acyl-CoA binding pocket adopts an open conformation in the MCC-apo structure (**D**) but switches to a close conformation in the MCC-ACO structure (**E**). (**F**) Alignment of the CT-N subdomain in the MCC-ACO structure with that in the PaMCC structure (PDB code: 3U9S)(*10*). The covalently linked biotin and its binding pocket adopt similar conformation in the two structures.

We aligned the MCC-ACO structure with the crystal structure of a bacteria MCC (PaMCC) holoenzyme in complex with CoA (PDB code: 3U9S) and observed a high similarity between them(*10*). Both the BCCP domain and the CT domain in the MCC-ACO structure align well with those in the PaMCC structure. The AMKM motif and biotin in the two structures also show a high similarity (Fig. 4F). Interestingly, alignment of the CoA-bound PaMCC holoenzyme structure with the structure of the PaMCC holoenzyme alone shows a change in the conformation of two helices in the CT-C subdomain of PaMCC(*10*), similar to that observed in the alignment of the MCC-ACO structure with the MCC-apo structure (Fig. 4C). These findings suggest that conformational changes in the acyl-CoA binding pocket, combined with the change in the position of the BCCP domain, may be necessary for the catalytic process of MCC.

## Discussion

Several three-dimensional structures of PCCs and MCCs have been reported(*9, 10, 15, 16, 18–20*), however, most of them are from bacteria and expressed in *Escherichia coli*(*9, 10, 15, 16, 18*). In our study, we extracted the endogenous PCC and MCC from human cells by one-step affinity purification (fig. S1) and determined their cryo-EM structures in the apo and substrate-bound states. The overall structures of human PCC and MCC holoenzymes resemble those of their bacterial homologs. This similarity aligns with the conserved functions of PCCs and MCCs across species, ranging from prokaryotes to eukaryotes.

A previous study showed that PCC purified from human liver also catalyzed the carboxylation of acetyl-CoA, but the rate was only about 1.5% of that observed for propionyl-CoA(*21*). Propionyl-CoA and acetyl-CoA, which have highly similar structures, undergo carboxylation at analogous positions—the α carbon of the propionyl or the acetyl group. A key question is, what determines the substrate specificity of the PCC holoenzyme? In the cryo-EM maps of the PCC holoenzymes, the acyl groups of acetyl-CoA and propionyl-CoA were not resolved (fig. S6), limiting the analysis of the interactions between the acyl groups and PCC. Nevertheless, the PCC-PCO and PCC-ACO structures determined in our study demonstrate that the conformations of the acyl-CoA binding pockets in the two structures are almost identical (Fig. 3F, fig. S7, B and C). In addition, the well resolved CoA groups of propionyl-CoA and acetyl-CoA bind at the same position in human PCC holoenzyme (Fig. 3F). These findings indicate that propionyl-CoA and acetyl-CoA bind to PCC with a similar binding mode. Thus, the selectivity of PCC towards propionyl-CoA and acetyl-CoA is largely determined by the interactions between the acyl group and PCC. The longer acyl group in propionyl-CoA, in comparison to that in acetyl-CoA, may mediate a stronger hydrophobic interaction that stabilizes the α carbon of the acyl group at a proper position to react with the carboxylated biotin.

The cryo-EM structures of MCC-apo and MCC-ACO reveal that there are two distinct binding sites in the CT domains of human MCC holoenzyme for the covalently linked biotin (Fig. 4B). In the MCC-apo structure, the biotin binds to a site that is distant from the catalytic residues A447 and G448, thus referred to as the exo-site; while in the MCC-ACO structure, the biotin binds to a site that is closer to A447 and G448 and is therefore referred to as the endo-site (Fig. 4B, fig. S10). The acyl-CoA binding pocket of the CT domains shows a different conformation in the MCC-ACO structure compared to that in the MCC-apo structure, and the covalently linked biotin is relocated from the exo-site to the endo-site. In the crystal structures of PaMCC in complex with CoA, the CT domains also adopt a different conformation in comparison with that in the apo structure of PaMCC(*10*). These findings indicate a coordination between biotin binding and acetyl-CoA binding for the MCC holoenzymes.

In another study from our group(*17*), it was also observed that the covalently linked biotin in the ACC-citrate filament, which is the active form of human ACC, binds to an exo-site that is distant from the acetyl-CoA binding site in the absence of acetyl-CoA. In contrast, in the crystal structure of CoA-bound yeast ACC, the biotin occupies an endo-site that is closer to the acetyl-CoA binding site(*22*).

In the crystal structure of *Staphylococcus aureus* PYC (SaPYC), two distinct binding sites for the covalently linked biotin were also observed, one is about 20 Å from the active site of the CT domain and is called the exo site, and the other is in the active site of the CT domain(*1, 23*).

Taken together, our studies, combined with previous research, suggest that the presence of an exo-site and an endo-site for the binding of covalently linked biotin is a common feature shared by many BDCs. The transfer of biotin from the exo-site to the endo-site may be promoted by the coordinated binding of acyl-CoA or pyruvate to the CT domains. Conversely, relocation of biotin from the endo-site to the exo-site might facilitate the release of the products after the second step carboxylation reaction.

However, we did not observe the relocation of biotin to an endo-site upon acyl-CoA binding in human PCC holoenzyme (Fig. 3). Additionally, while we observed differences in the binding position of biotin and the conformation of the acyl-CoA binding pocket in the MCC-ACO structure compared to those in the MCC-apo structure, such differences were not observed in the MCC-PCO structure. It is noteworthy that neither acetyl-CoA nor propionyl-CoA is the natural substrate of MCC. Recently, a cryo-EM structure of the human MCC holoenzyme in complex with its natural substrate, MC-CoA, was resolved (PDB code: 8J4Z). In this structure, the binding site of biotin and the conformation of the CT domains closely resemble that in our acetyl-CoA-bound MCC structure (Fig. 4E and fig. S11). Therefore, the binding of acetyl-CoA mimics the binding of MC-CoA. Further studies focusing on capturing the structures of human PCC and MCC holoenzymes in various states during their catalytic processes would contribute to a comprehensive understanding of the underlying catalytic mechanisms.

## Materials and Methods

### Cell culture

Expi 293F cells (Thermo Scientific) were cultured in SMM 293T-II medium (Sino Biological) at 37 °C with 5% CO_2_ in a Multitron-Pro incubator (Infors). After the density reached 2.0×10^6^/mL, the cells were cultured for an additional 48 h, then harvested by centrifuging at 4,000 ×*g* (Fiberlite F9-6×1000 LEX rotor, Thermo Scientific) for 10 min and resuspended with HBS buffer (20 mM HEPES, pH 7.4, 150 mM NaCl). The cells were snap frozen and stored at −80 °C.

### Endogenous purification of biotin dependent carboxylases

The frozen cells were thawed and disrupted using a high-pressure cell crusher and centrifuged at 23,376 ×*g* (JA-14.50 rotor, Beckman) for 1 h at 4 °C. The supernatant was incubated with Strep-Tactin^®^XT resin (IBA Lifesciences) for 1 h at 4 °C and then washed with HBS buffer. The proteins were eluted by HBS buffer supplemented with 50 mM-biotin (J&K Scientific) and further purified by size-exclusion chromatography (Superose 6 Increase 10/300 GL column, Cytiva) using HBS as the running buffer. Different batches of proteins were purified for the preparation of the samples at different substrate binding states. For the substrate-free sample (the apo sample), fractions eluted within 12.5–15 mL were combined and concentrated to 2.85 mg/mL. For the substrate-bound samples, fractions eluted within 12.5–16.5 mL were combined and concentrated to 2.5–2.65 mg/mL. The samples were pre-incubated with 25 mM NaHCO_3_, 10 mM MgCl_2_ and 10 mM propionyl-CoA for the propionyl-CoA-bound sample (the PCO sample) or acetyl-CoA for the acetyl-CoA-bound sample (the ACO sample), followed by adding 10 mM ATP immediately before frozen on the cryo-EM grids. The proteins were analyzed by mass spectrometry and applied for cryo-EM grids preparation the same day purified.

### Cryo-EM data acquisition and pre-processing

Holey grids (Holey carbon filmed 300 mesh gold grids, R1.2/1.3, Quantifoil) were glow discharged with a medium RF power for 30 s using a plasma cleaner PDC-32G-2 (Harrick Plasma) prior use. Aliquots (4 μL) of each sample were loaded to the grids and flash frozen in liquid ethane immediately using Vitrobot Mark IV (Thermo Fisher Scientific). The cryo-EM grids were stored in liquid nitrogen until data acquisition.

The cryo-EM grids were transferred to a Titan Krios TEM (FEI) operating at 300 kV equipped with a K3 Summit direct electron detector (Gatan) and a GIF Quantum energy filter (Gatan). Zero-loss movie stacks were automatically collected using EPU software (FEI) with a slit width of 20 eV on the energy filter and a defocus range from −1.2 to −2.2 µm in super-resolution mode at an 81,000× nominal magnification. Each movie stack, which contained 32 frames, was exposed for 2.56 s with a total electron dose of ∼50 e^-^/Å^2^. The stacks were motion corrected using MotionCor2 with a binning factor of 2, resulting in the pixel size of 1.0773 Å(*24*). Dose weighting was performed concurrently(*25*). The movie stacks were imported to cryoSPARC (v3.3.2, Structura Biotechnology) for CTF estimation and downstream processing(*26*).

### Data processing

For the apo sample, 11,481 micrographs were captured and imported to cryoSPARC for processing. Particles were automatically picked according to the templates generated from an initial 2D classification of the manually picked particles and extracted with a box size of 488 pixels for *ab-initio* reconstruction and heterogeneous refinement jobs, yielding the maps of PCC (11.5%) and MCC (37.6%) holoenzymes. To get the high-resolution map of PCC holoenzyme, a non-uniform refinement with the corresponding group of particles was carried out, yielding a detailed map, yet the density for one of the α subunits was lacking. To solve this problem, distinct reference volumes with (Ref-A) or without (Ref-B) the corresponding PCCα were generated by UCSF Chimera (v1.16)(*27*). After a refinement based on multiple references, 61,567 particles were subjected to the homogenous refinement, yielding the complete map of PCC (PCC-apo) at 3.02 Å resolution. For the case of MCC holoenzyme, a round of non-uniform refinement was carried out. After that, the volume chirality was flipped, and the map was further polished by a series of refinement jobs, resulting in the map of MCC (MCC-apo) at 2.29 Å resolution.

For the PCO sample, 10,898 micrographs were imported for data processing. After particle picking, extraction (box size = 488 pixels), *ab-initio* reconstruction and heterogeneous refinement, the PCC (8.7%) and MCC (39.7%) representing volumes were further processed by non-uniform refinement, yielding the maps generated from 105,293 and 476,773 particles, respectively. The PCC map was subjected to local CTF refinement and subsequent local refinement, yielding the final map of propionyl-CoA bound PCC holoenzyme (PCC-PCO) at 2.80 Å. The MCC map was further polished by heterogenous refinement and non-uniform refinement after flipping chirality, producing the final map of propionyl-CoA bound MCC holoenzyme (MCC-PCO) at 2.36 Å.

For the ACO sample, 4,036 micrographs were collected and imported. Particle picking, extraction (box size = 488 pixels), initial 2D classification and particle clustering jobs were carried out accordingly. 31,622 PCC-like particles were enriched in cluster-A and subsequently subjected to *ab-initio* reconstruction and non-uniform refinement. After flipping the volume chirality, the refined map was further polished by another round of non-uniform refinement, yielding the 3.38 Å map of acetyl-CoA bound PCC holoenzyme (PCC-ACO). In cluster-B, 273,428 MCC-like particles were enriched. These particles were *ab-initio* reconstructed and classified to 4 volumes, of which one, composed of 39.9% of particles, exhibited clear secondary structure features of MCC. This map was further processed by heterogeneous refinement and non-uniform refinement jobs, yielding the final map of acetyl-CoA bound MCC holoenzyme (MCC-ACO) at 2.85 Å resolution.

The resolutions mentioned above were determined according to the gold-standard Fourier shell correlation 0.143 criterion and with a high-resolution noise substitution method(*28*). All the reconstructed maps were subjected to further model building.

### Model building and structure refinement

The AlphaFold predicted structures were used as the templates for the model building of PCC and MCC holoenzymes(*29, 30*). The reference models were automatically placed in the EM maps using Rosetta and manually adjusted using Coot, resulting the modified models(*31, 32*). Each residue was manually checked with the chemical properties taken into consideration during model building. Several segments, whose corresponding densities were invisible, were not modeled. Real space refinement was performed using Phenix with secondary structure and geometry restraints to prevent overfitting(*33*). For the cross-validation of the structure, the model was refined against one of the two independent half maps from the gold-standard 3D refinement approach. Then, the refined model was tested against the other half map. Statistics associated with data collection, 3D reconstruction and model building were summarized in tables S1 and S2.

## Acknowledgments

We thank Drs. Xiaofeng Zhang and Xiechao Zhan for valuable discussion and suggestions on structure determination. We thank the Cryo-EM Facility, the High-Performance Computing Center, and the Mass Spectrometry & Metabolomics Core Facility of Westlake University for technical support.

## Author contributions

Conceptualization: Q.H., F.Z.

Methodology: Q.H., F.Z.

Investigation: F.Z., Y.Zhang, Y.Zhu

Visualization: F.Z., Y.Zhang

Funding acquisition: Q.H.

Project administration: Q.H., F.Z.

Supervision: Q.H., Y.S.

Writing – original draft: Q.H., F.Z.

Writing – review & editing: Q.H., Y.S., Q.Z.

## Data and materials availability

The cryo-EM structures have been deposited in the Protein Data Bank with the accession codes 8XL3 (PCC-apo), 8XL4 (PCC-ACO), 8XL5 (PCC-PCO), 8XL6 (MCC-apo), 8XL7 (MCC-ACO) and 8XL8 (MCC-PCO). The cryo-EM maps have been deposited in the Electron Microscopy Data Bank with the accession codes EMD-38436 (PCC-apo), EMD-38437 (PCC-ACO), EMD-38438 (PCC-PCO), EMD-38439 (MCC-apo), EMD-38440 (MCC-ACO) and EMD-38441 (MCC-PCO). The predicted structures of human PCC and MCC subunits used for cryo-EM model building were downloaded from the AlphaFold Protein Structure Database with the accession codes AF-P05165-F1 (PCCα), AF-P05166-F1 (PCCβ), AF-Q96RQ3-F1 (MCCα) and AF-Q9HCC0-F1 (MCCβ). The structures of bacteria PCC (PDB codes: 1XNY, 3N6R, 6YBP), bacteria MCC (PDB codes: 3U9S and 3U9T), and 3-methylcrotonyl-CoA-bound human MCC (PDB code: 8J4Z) were used for structural analysis in this study.

## Supplementary Materials

**Fig. S1.**
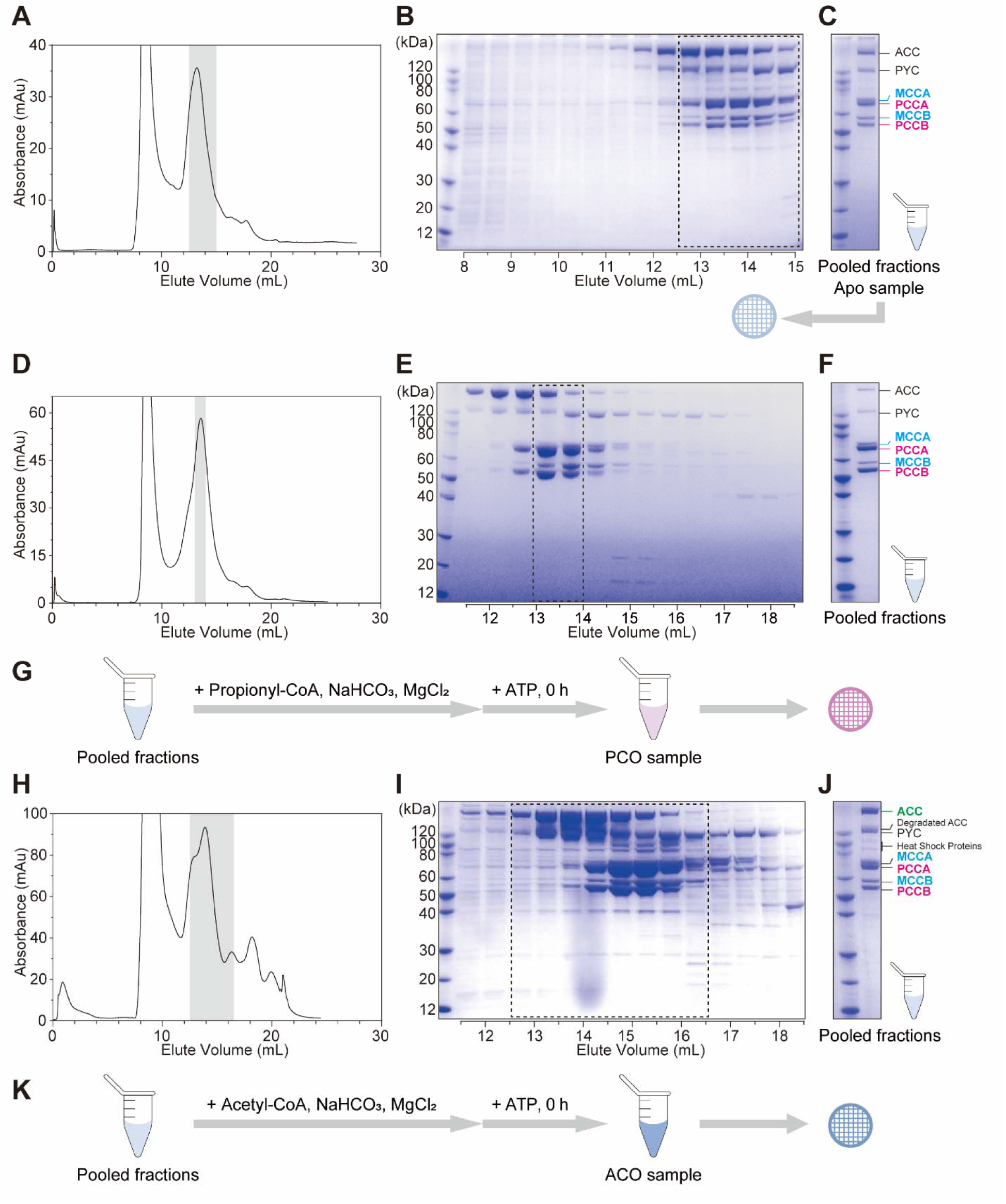
Protein purification and cryo-EM sample preparation. (**A**–**C**) Preparation of the apo sample. The proteins eluted from Strep-Tactin^®^XT resin was further purified by size-exclusion chromatography (**A**). The fractions were analyzed by SDS-PAGE and visualized by Coomassie blue staining (**B**). The peak fractions were combined and analyzed by mass spectrometry, and the concentrated sample was directly frozen on a Holey Carbon filmed 300-mesh gold grid (Quantifoil) (**C**). (**D**–**G**) Preparation of the PCO sample. The proteins eluted from Strep-Tactin^®^XT resin was further purified by size-exclusion chromatography (**D**). The fractions were analyzed by SDS-PAGE and visualized by Coomassie blue staining (**E**). The peak fractions were combined and analyzed by mass spectrometry (**F**). The purified proteins were concentrated and incubated with propionyl-CoA, NaHCO_3_ and MgCl_2_, and mixed with ATP before frozen on a Holey Carbon filmed 300-mesh gold grid (Quantifoil) (**G**). (**H**–**K**) Preparation of the ACO sample. The proteins eluted from Strep-Tactin^®^XT resin was further purified by size-exclusion chromatography (**H**). The fractions were analyzed by SDS-PAGE and visualized by Coomassie blue staining (**I**). The peak fractions were combined and analyzed by mass spectrometry (**J**). The purified proteins were concentrated and incubated with acetyl-CoA, NaHCO_3_ and MgCl_2_, and mixed with ATP before frozen on a Holey Carbon filmed 300-mesh gold grid (Quantifoil) (**K**).

**Fig. S2.**
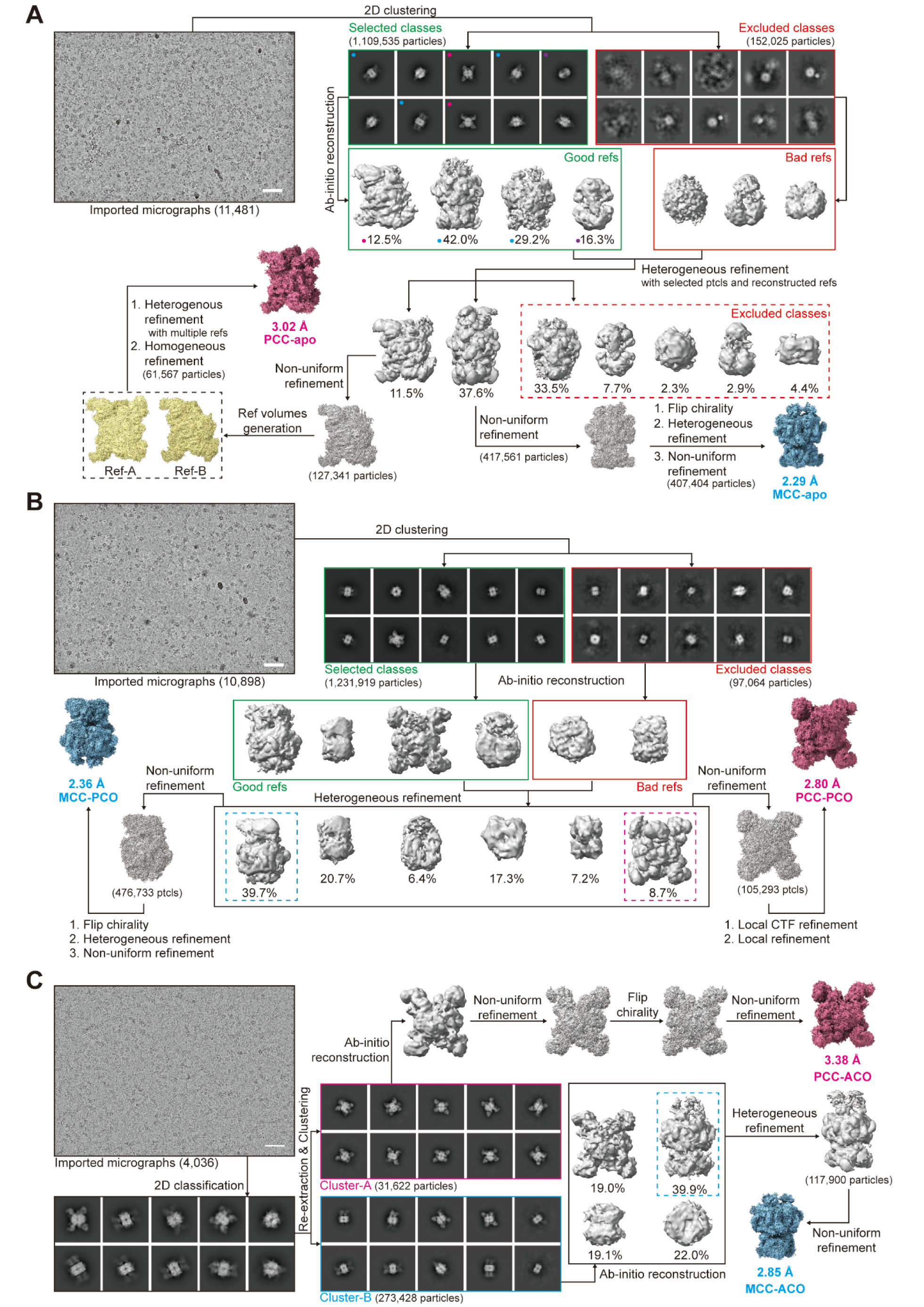
Flowchart for the cryo-EM data processing of human PCC and MCC holoenzymes. (**A**) Cryo-EM data processing of the PCC-apo and MCC-apo structures. Particles from 11,481 micrographs went through multiple rounds of classification to reconstruct several cryo-EM density maps. The final cryo-EM density maps achieved the resolutions of 3.02 Å and 2.29 Å for PCC-apo and MCC-apo, respectively. For the 2D classes and 3D volumes, the red dots indicate PCC, and the blue dots indicate MCC. (**B**) Cryo-EM data processing of the PCC-PCO and MCC-PCO structures. Particles from 10,898 micrographs went through multiple rounds of classification to reconstruct several cryo-EM density maps. The final cryo-EM density maps achieved the resolutions of 2.80 Å and 2.36 Å for PCC-PCO and MCC-PCO, respectively. (**C**) Cryo-EM data processing of the PCC-ACO and MCC-ACO structures. Particles from 4,036 micrographs went through multiple rounds of classification to reconstruct several cryo-EM density maps. The final cryo-EM density maps achieved the resolutions of 3.38 Å and 2.85 Å for PCC-ACO and MCC-ACO, respectively. The scale bar in each micrograph is 50 nm.

**Fig. S3.**
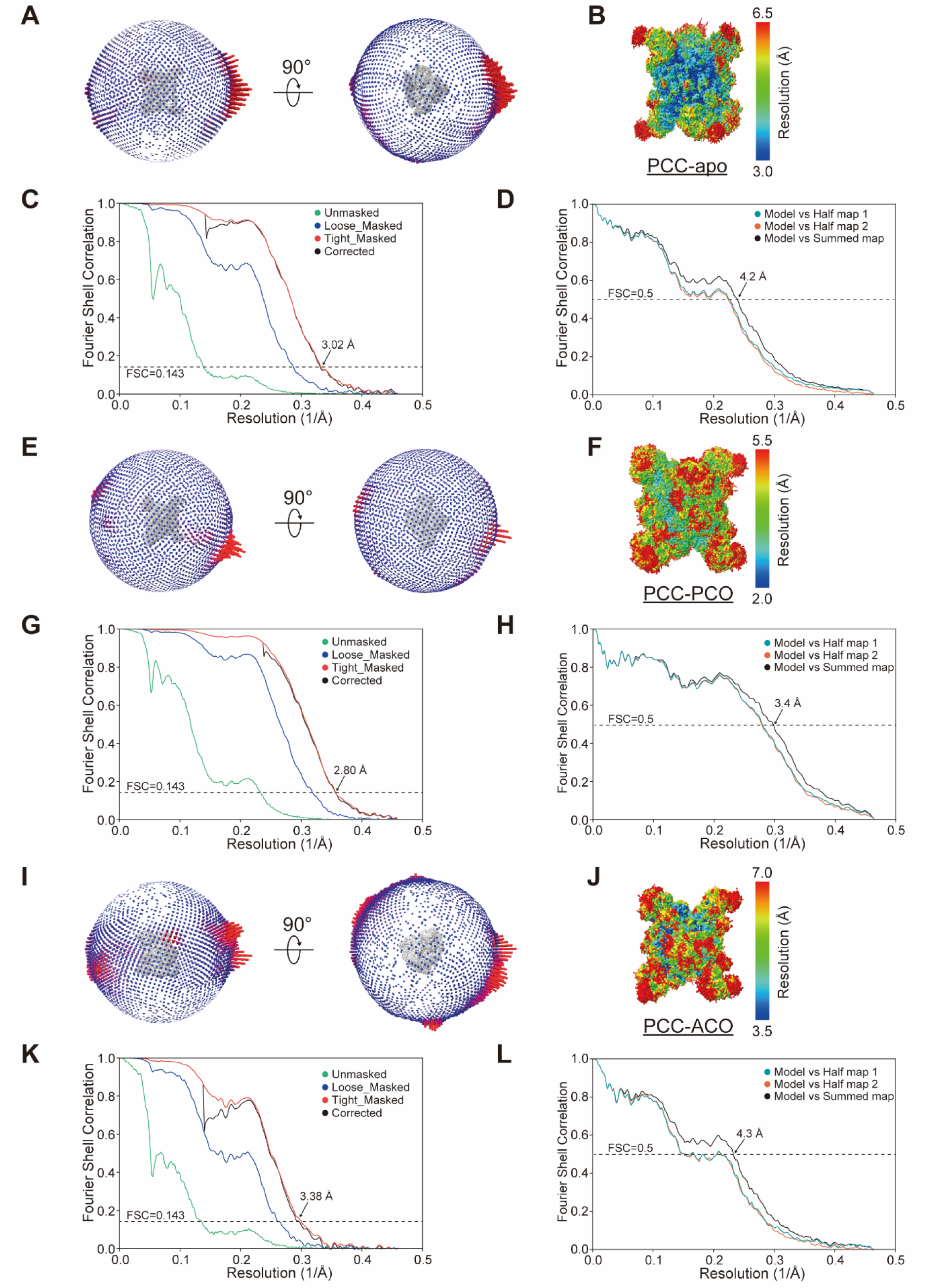
Cryo-EM analysis of human PCC holoenzymes. (**A**–**D**) Cryo-EM analysis of the PCC-apo structure. (**A**) Cryo-EM map and the corresponding Euler distribution plot of the PCC-apo structure. (**B**) The local resolution estimation of the PCC-apo structure. (**C**) The gold standard FSC curves and (**D**) the FSC curves for cross-validation of the PCC-apo structure. (**E**–**H**) Cryo-EM analysis of the PCC-PCO structure. (**E**) Cryo-EM map and the corresponding Euler distribution plot of the PCC-PCO structure. (**F**) The local resolution estimation of the PCC-PCO structure. (**G**) The gold standard FSC curves and (**H**) the FSC curves for cross-validation of the PCC-PCO structure. (**I**–**L**) Cryo-EM analysis of the PCC-ACO structure. (**I**) Cryo-EM map and the corresponding Euler distribution plot of the PCC-ACO structure. (**J**) The local resolution estimation of the PCC-ACO structure. (**K**) The gold standard FSC curves and (**L**) the FSC curves for cross-validation of the PCC-ACO structure. The particles were extracted with a box size of 488 pixels. The map resolution was determined by the reciprocal of the spatial frequency at FSC = 0.143, and the consistency of the structure optimization process was verified by comparison of the molecular model with the summed (the black curves) or half (the red or green curves) maps at FSC = 0.5(*28, 33*).

**Fig. S4.**
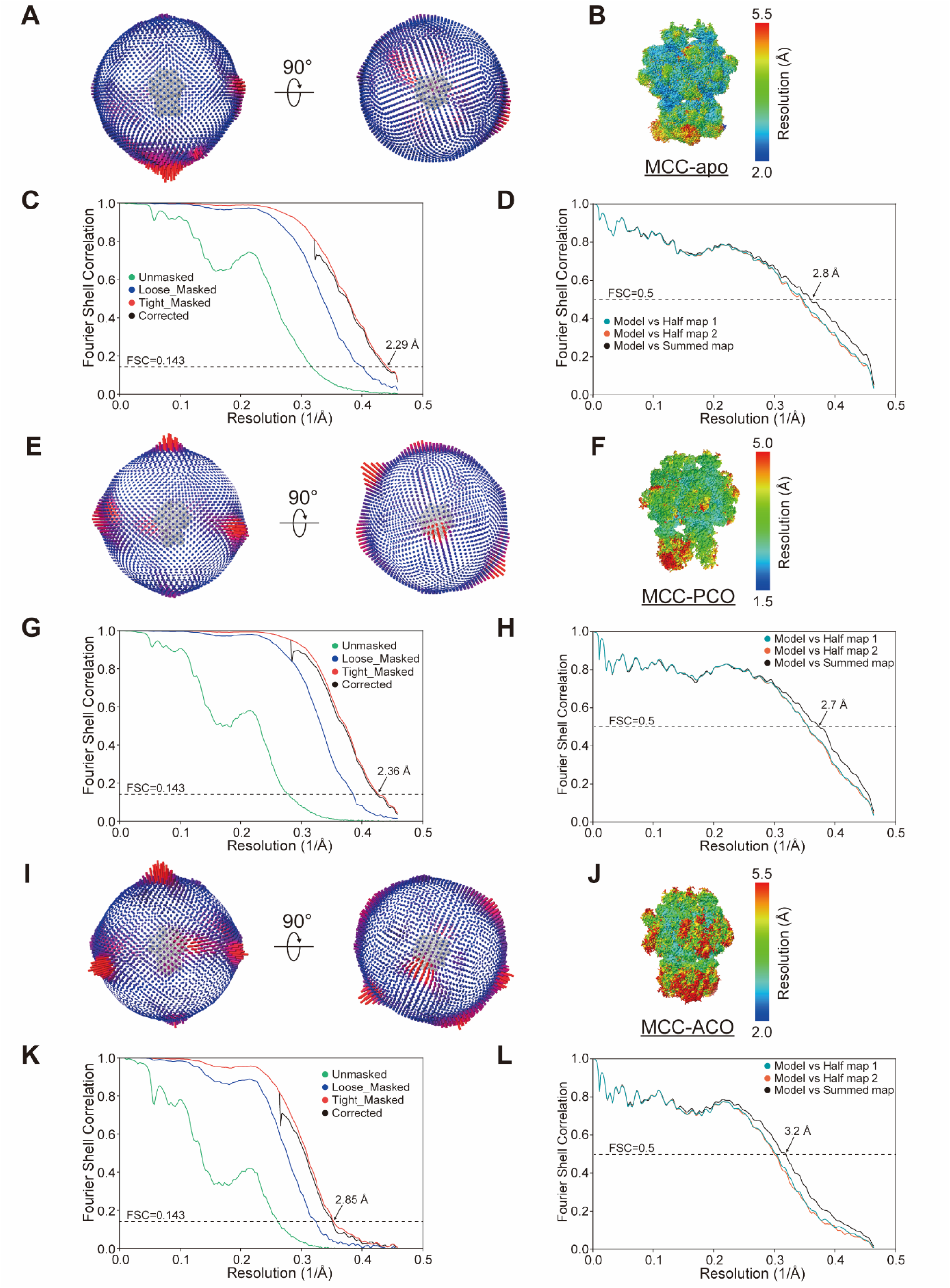
Cryo-EM analysis of human MCC holoenzymes. (**A**–**D**) Cryo-EM analysis of the MCC-apo structure. (**A**) Cryo-EM map and the corresponding Euler distribution plot of the MCC-apo structure. (**B**) The local resolution estimation of the MCC-apo structure. (**C**) The gold standard FSC curves and (**D**) the FSC curves for cross-validation of the MCC-apo structure. (**E**– **H**) Cryo-EM analysis of the MCC-PCO structure. (**E**) Cryo-EM map and the corresponding Euler distribution plot of the MCC-PCO structure. (**F**) The local resolution estimation of the MCC-PCO structure. (**G**) The gold standard FSC curves and (**H**) the FSC curves for cross-validation of the MCC-PCO structure. (**I**–**L**) Cryo-EM analysis of the MCC-ACO structure. (**I**) Cryo-EM map and the corresponding Euler distribution plot of the MCC-ACO structure. (**J**) The local resolution estimation of the MCC-ACO structure. (**K**) The gold standard FSC curves and (**L**) the FSC curves for cross-validation of the MCC-ACO structure. The particles were extracted with a box size of 488 pixels. The map resolution was determined by the reciprocal of the spatial frequency at FSC = 0.143, and the consistency of the structure optimization process was verified by comparison of the molecular model with the summed (the black curves) or half (the red or green curves) maps at FSC = 0.5(*28, 33*).

**Fig. S5.**
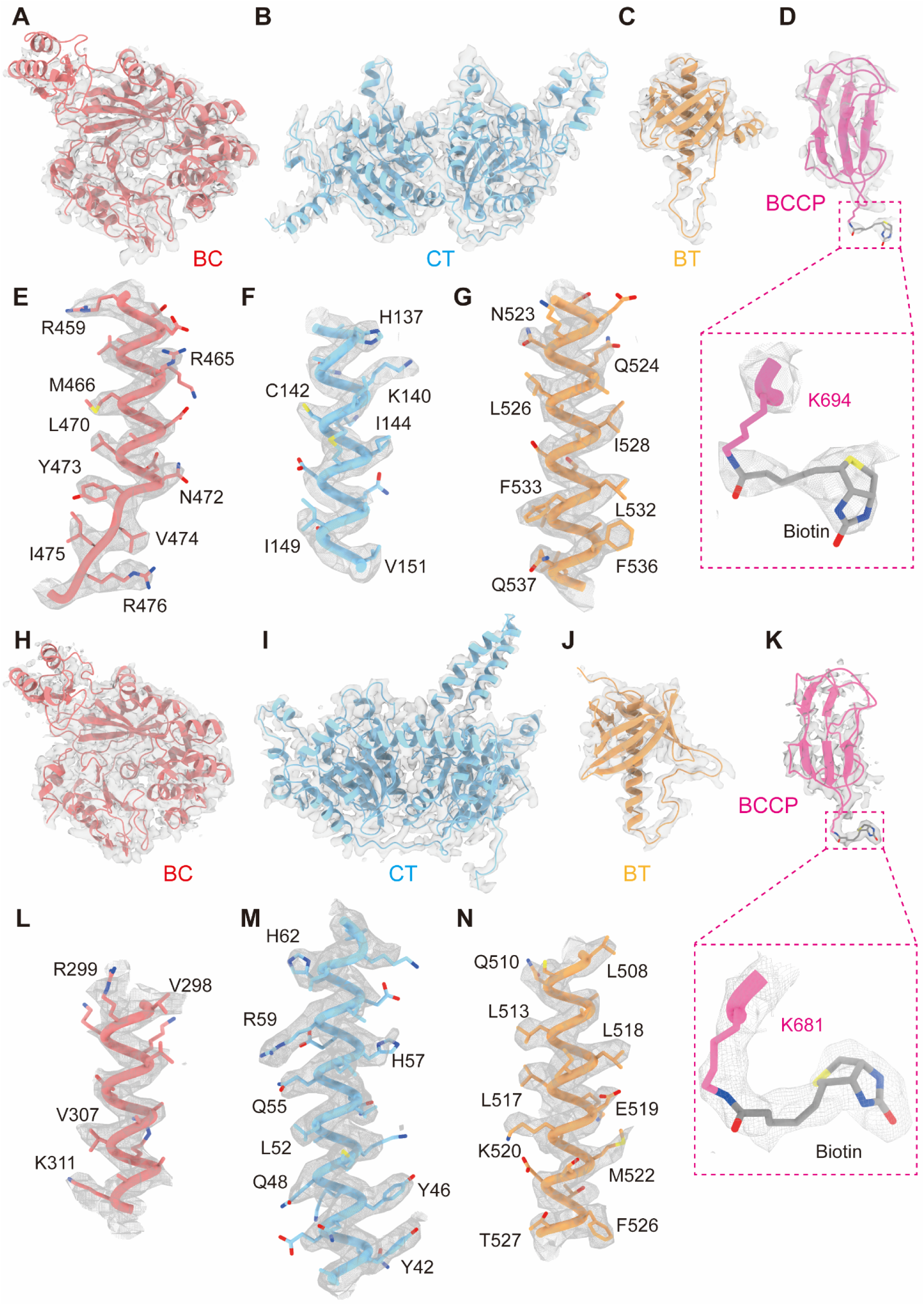
The local density of human PCC and MCC holoenzymes. (**A**–**D**) Cryo-EM maps of the BC domain (**A**), CT domain (**B**), BT domain (**C**) and biotinylated BCCP domain (**D**) in the PCC-apo structure. The biotin is covalently linked to K694 at the AMKM motif of the BCCP domain (**D**). (**E**–**G**) Cryo-EM maps of representative α-helices in the BC (**E**), CT (**F**) and BT (**G**) domains in the PCC-apo structure. (**H**–**K**) Cryo-EM maps of the BC domain (**H**), CT domain (**I**), BT domain (**J**) and biotinylated BCCP domain (**K**) in the MCC-apo structure. The biotin is covalently linked to K681 at the AMKM motif of the BCCP domain (**K**). (**L**–**N**) Cryo-EM maps of representative α-helices in the BC (**L**), CT (**M**) and BT (**N**) domains in the MCC-apo structure.

**Fig. S6.**
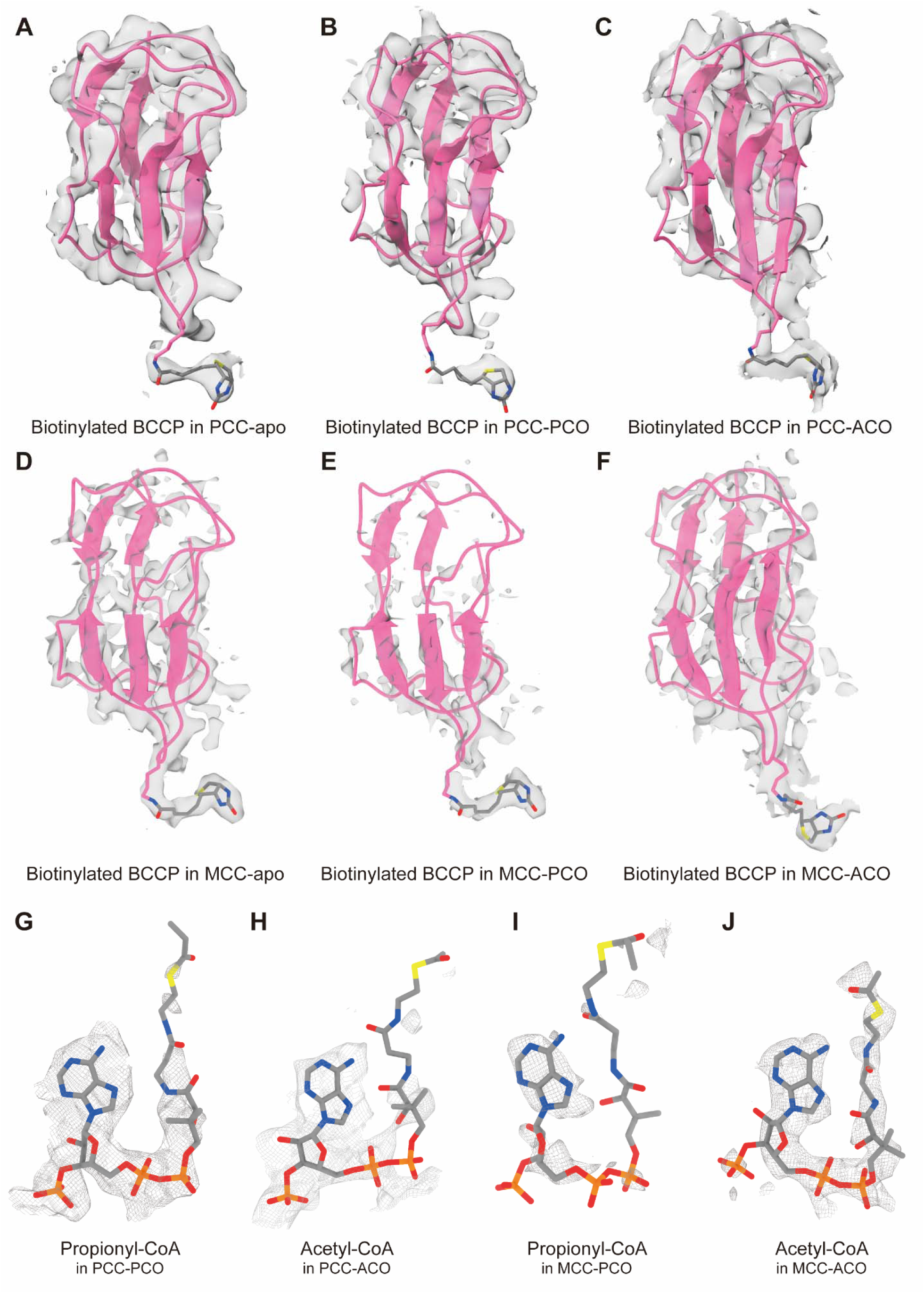
The local density of the biotinylated BCCP domains and acyl-CoA in human PCC and MCC holoenzymes. (**A**–**C**) Cryo-EM maps of the biotinylated BCCP in the PCC-apo (**A**), PCC-PCO (**B**) and PCC-ACO (**C**) structures. (**D**–**F**) Cryo-EM maps of the biotinylated BCCP in the MCC-apo (**D**), MCC-PCO (**E**) and MCC-ACO (**F**) structures. (**G**–**J**) Cryo-EM maps of the propionyl-CoA in the PCC-PCO structure (**G**), the acetyl-CoA in the PCC-ACO structure (**H**), the propionyl-CoA in the MCC-PCO structure (**I**) and the acetyl-CoA in the MCC-ACO structure (**J**). The cryo-EM maps were displayed at an RMS level of 5 σ for (**A**–**F**) and of 4.5 σ for (**G**–**J**).

**Fig. S7.**
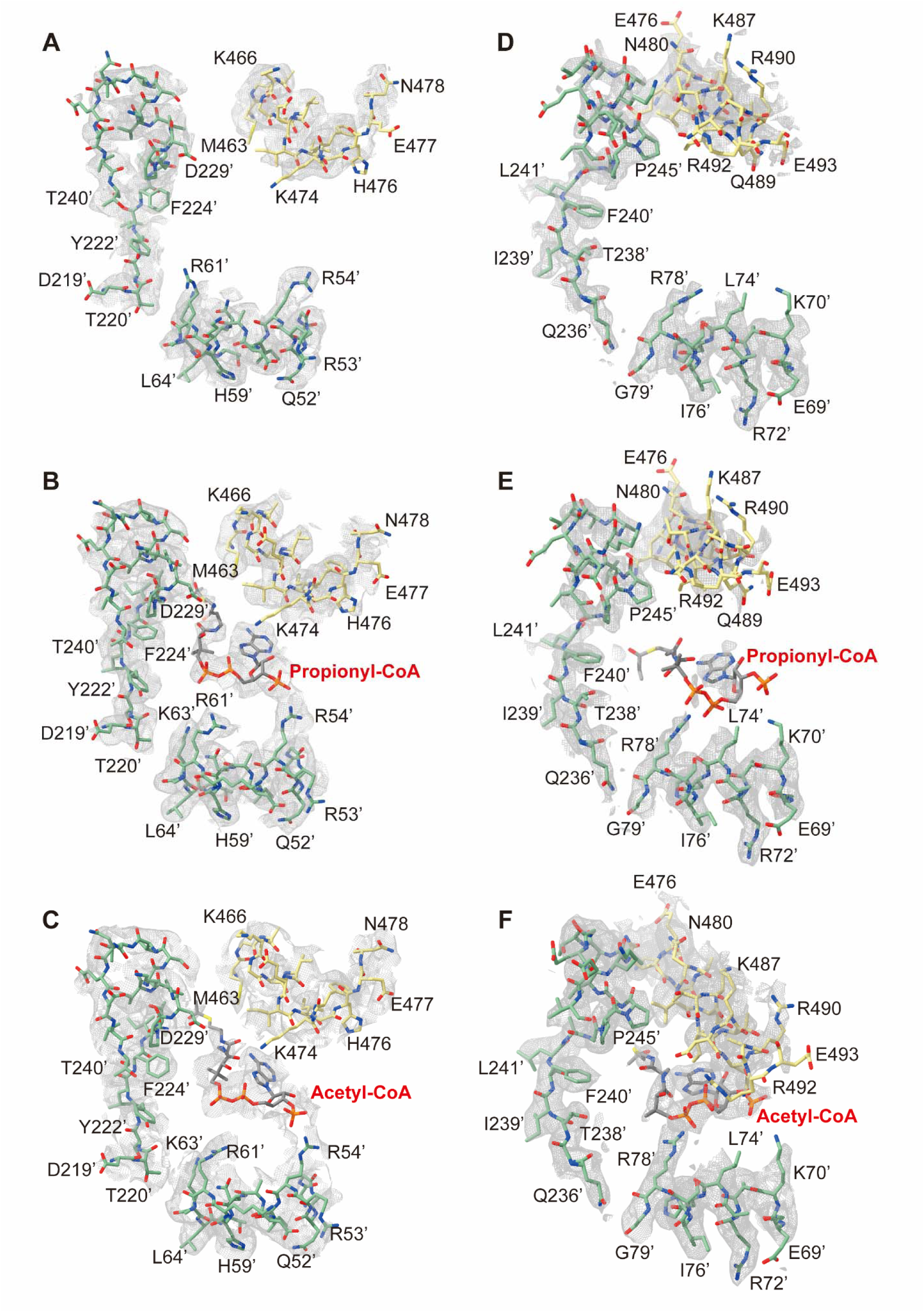
The local density of the acyl-CoA binding pocket in human PCC and MCC holoenzymes. (**A**–**C**) Cryo-EM maps of the acyl-CoA binding pocket in the PCC-apo (**A**), PCC-PCO (**B**) and PCC-ACO (**C**) structures. (**D**–**F**) Cryo-EM maps of the acyl-CoA binding pocket in the MCC-apo (**D**), PCC-PCO (**E**) and PCC-ACO (**F**) structures. All cryo-EM maps were displayed at an RMS level of 4.5 σ.

**Fig. S8.**
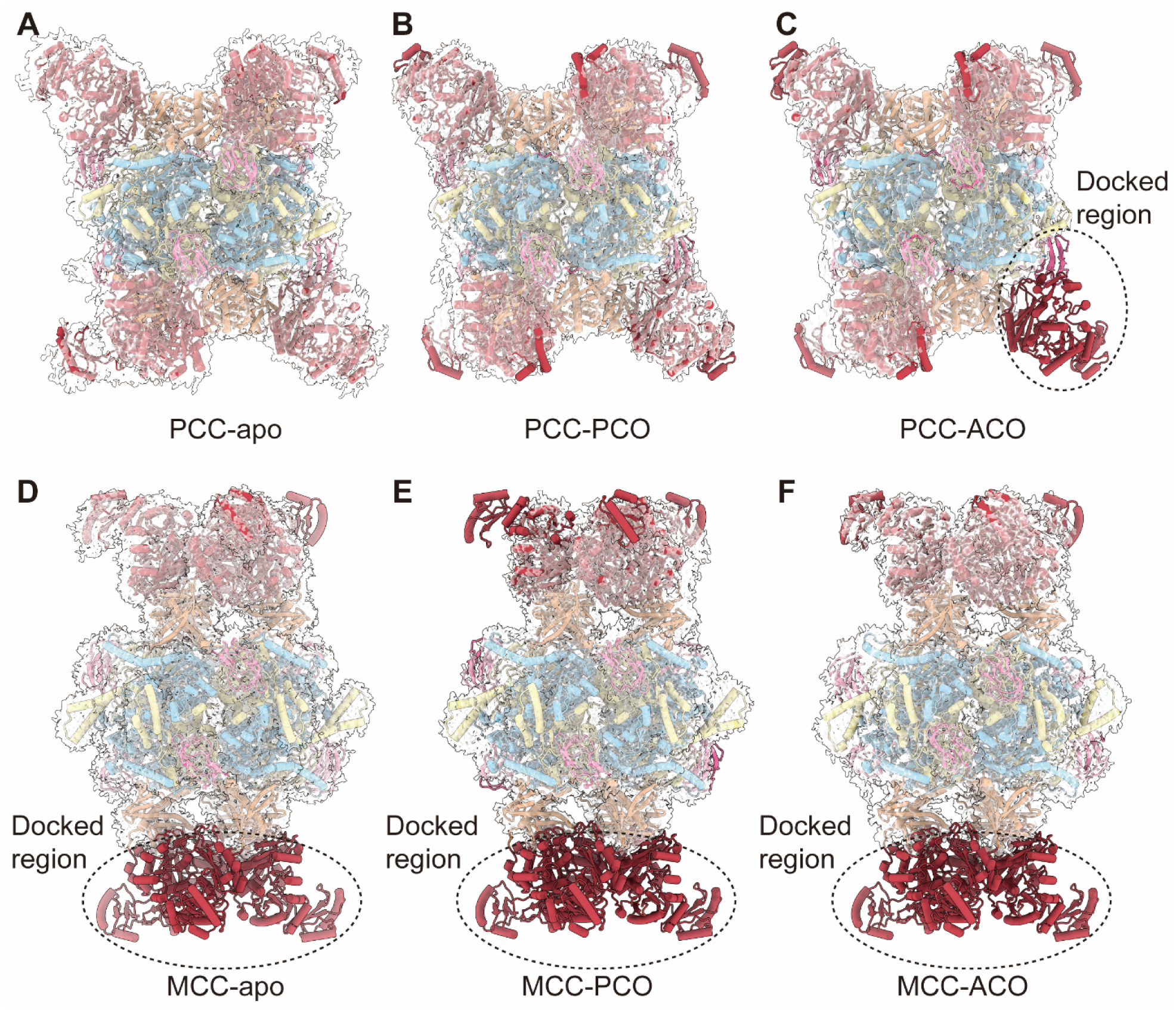
The Cryo-EM maps of human PCC and MCC holoenzymes. (**A**–**C**) Cryo-EM maps for the PCC-apo (RMS Level = 2.8 σ) (**A**), PCC-PCO (RMS Level = 2.7 σ) (**B**) and PCC-ACO (RMS Level = 3.5 σ) (**C**). The cryo-EM maps are displayed in semitransparent white color and superimposed with the corresponding stereo models. The density of one PCCα subunit in the PCC-ACO structure is partially missing (indicated by dashed circle in panel **C**). (**D**–**F**) Cryo-EM maps for the MCC-apo (RMS Level = 2.3 σ) (**D**), MCC-PCO (RMS Level = 2.0 σ) (**E**) and MCC-ACO (RMS Level= 2.8 σ) (**F**). The density of three MCCα subunits in each MCC holoenzyme structure are partially missing (indicated by dashed circles in panels **D**–**F**).

**Fig. S9.**
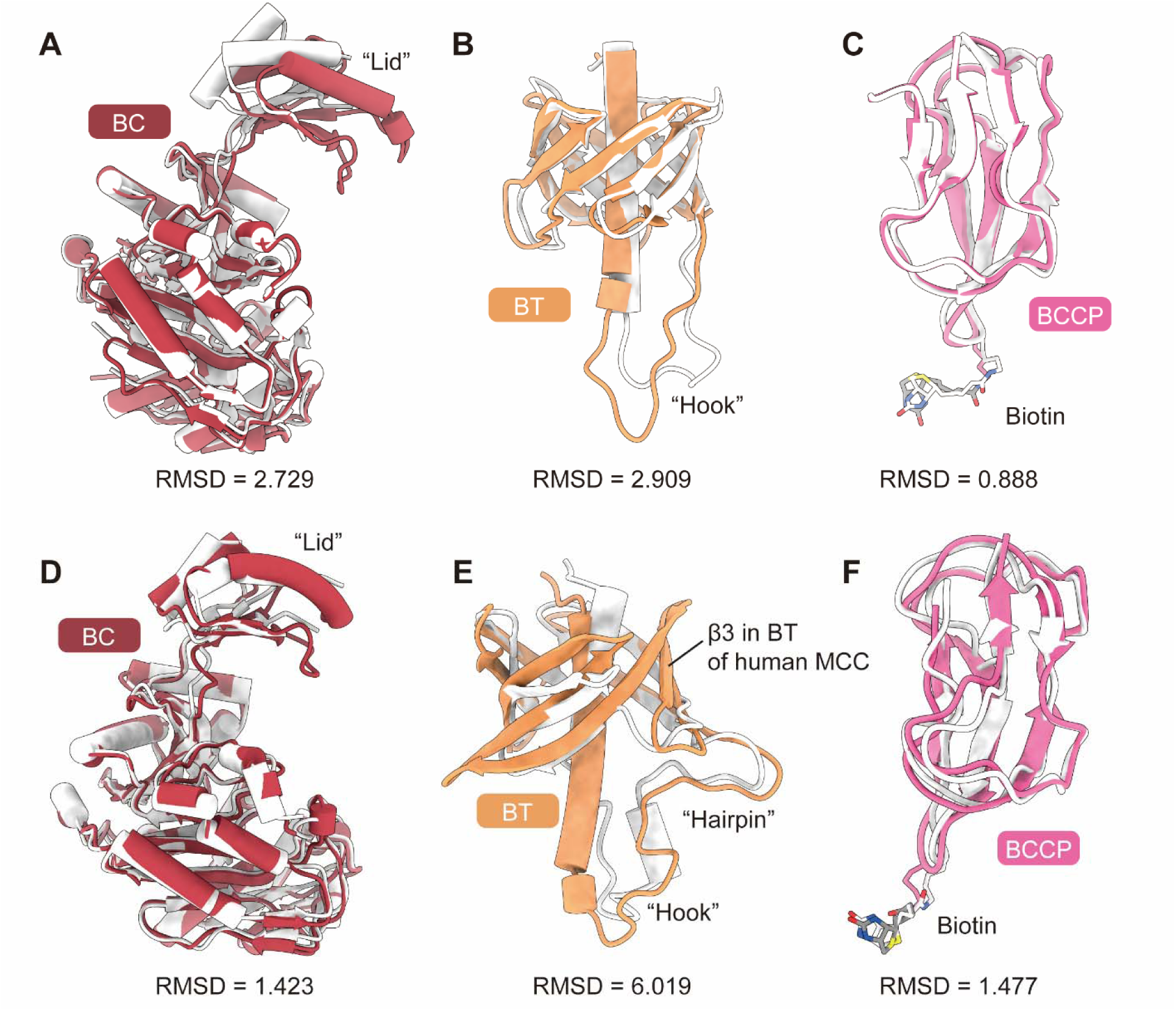
Comparison of the domain structures of human PCC and MCC with that of their bacteria homologs. (**A**–**C**) Alignments of the BC domain (**A**), BT domain (**B**) and biotinylated BCCP domain (**C**) in human PCC structure (PCC-apo, colored) with that in a bacteria PCC structure (PDB code: 3N6R, gray)(*9*). (**D**–**F**) Alignments of the BC domain (**D**), BT domain (**E**) and biotinylated BCCP domain (**F**) in human MCC structure (MCC-apo, colored) with that in a bacteria MCC structure (PDB code: 3U9T, gray)(*10*).

**Fig. S10.**
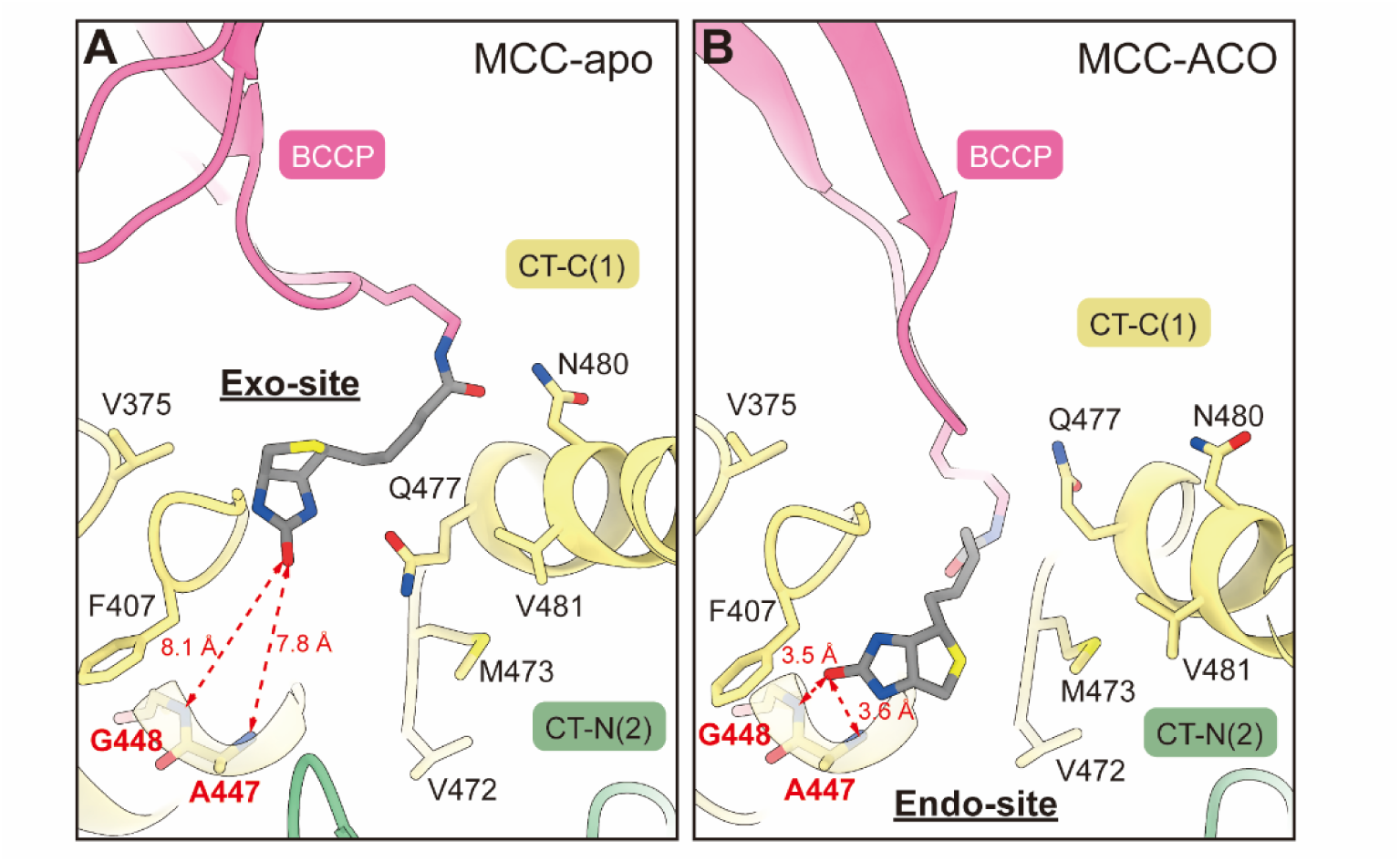
The biotin binding sites in the structures of human MCC holoenzyme. (**A**) The covalently linked biotin in the MCC-apo structure binds to a site distant from the catalytic residues in the CT domain. This site is thus referred to as the exo-site. The distances between the carbonyl oxygen of the ureido ring of biotin and the main chain nitrogen atoms of the catalytic residues A447 and G448 in the CT domain are 7.8 and 8.1, respectively. (**B**) The covalently linked biotin in the MCC-ACO structure moves to a site closer to the catalytic residues A447 and G448 in the CT domain. This site is referred to as the endo-site. The distances between the carbonyl oxygen of the ureido ring of biotin and the main chain nitrogen atoms of the catalytic residues A447 and G448 in the CT domain are shortened to 3.6 and 3.5, respectively.

**Fig. S11.**
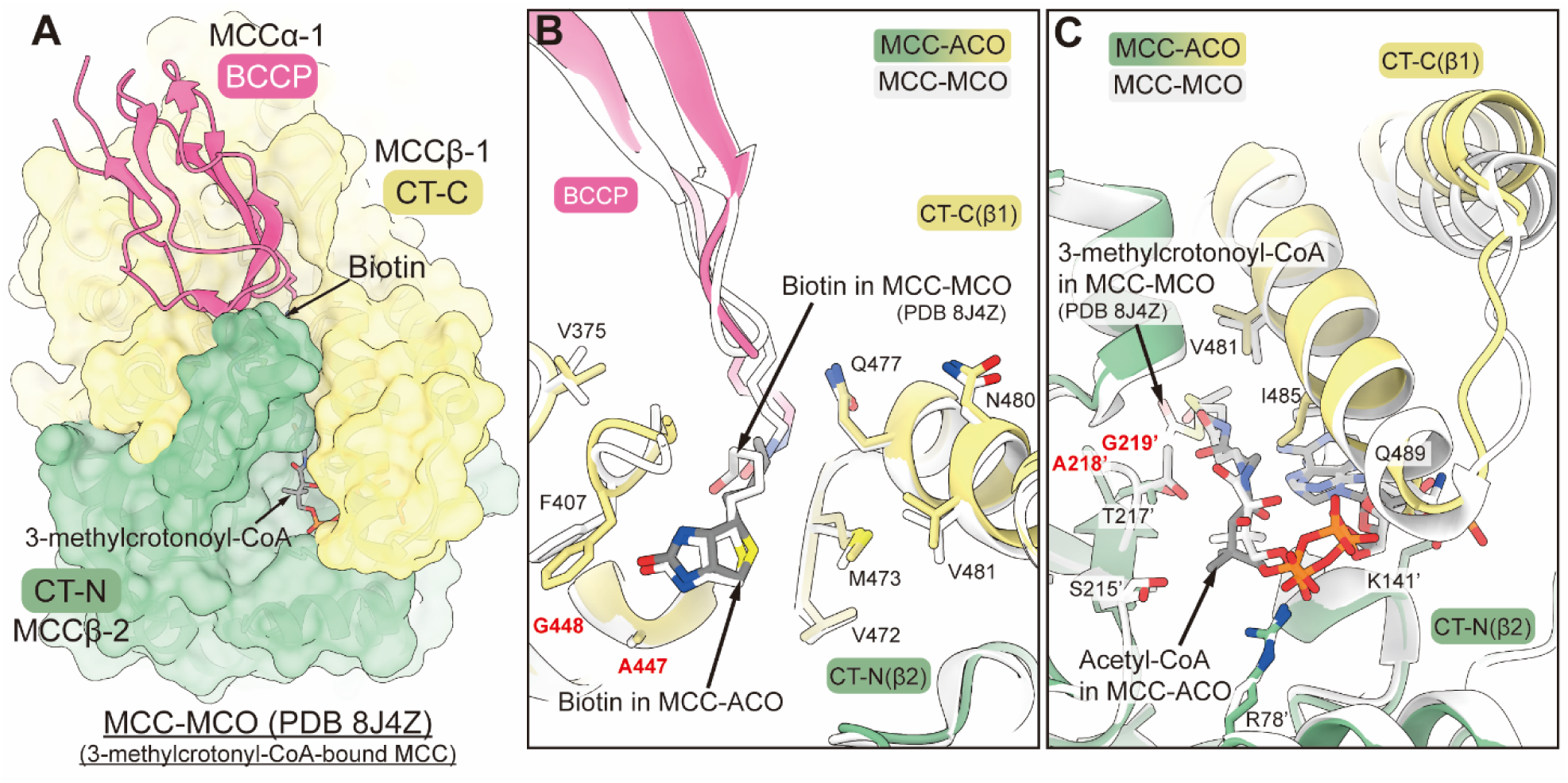
Comparison of human MCC holoenzyme structures in complex with acetyl-CoA and 3-methylcrotonyl-CoA. (**A**) Surface representation of the binding pocket of biotin and acyl-CoA in human MCC holoenzyme structure in complex with its natural substrate, 3-methylcrotonyl-CoA (PDB code: 8J4Z). The acyl-CoA binding pocket adopts a closed conformation, which is similar to that in the structure of acetyl-CoA-bound MCC (refer to Fig. 4E). (**B**) Alignment of the CT-N subdomain in the MCC-ACO structure with that in the 3-methylcrotonyl-CoA-bound MCC (MCC-MCO) structure. The biotin molecules in both structures bind to an endo-site, and the biotin binding pockets adopt a similar conformation. (**C**) Alignment of the CT-N subunit in the MCC-ACO structure with that in the 3-methylcrotonyl-CoA-bound MCC (MCC-MCO) structure.

**Table S1.** Cryo-EM data collection, reconstruction, refinement, and validation statistics of human PCC structures.

| Sample | PCC-apo | PCC-ACO | PCC-PCO |
| --- | --- | --- | --- |
| PDB code | 8XL3 | 8XL4 | 8XL5 |
| EMDB code | EMD-38436 | EMD-38437 | EMD-38438 |
| Data collection |  |  |  |
| EM equipment | Titan Krios (Thermo Fisher Scientific) |  |  |
| Voltage (kV) | 300 |  |  |
| Detector | Gatan K3 Summit |  |  |
| Energy filter | Gatan GIF Quantum, 20 eV slit |  |  |
| Pixel size (Å) | 1.0773 |  |  |
| Electron dose (e<sup>-</sup>/Å<sup>2</sup>) | 50 |  |  |
| Defocus range (μm) | −1.2 to −2.2 |  |  |
| Number of collected micrographs | 11,481 | 4,036 | 10,898 |
| 3D Reconstruction |  |  |  |
| SoftwaresSoftware | cryoSPARC 3.3.2 |  |  |
| Number of used particles | 61,567 | 30,932 | 105,293 |
| Resolution (Å) | 3.02 | 3.38 | 2.80 |
| Symmetry | C1 |  |  |
| Map sharpening B-factor (Å<sup>2</sup>) | 68 | 70.3 | 75.8 |
| Refinement |  |  |  |
| Software | Phenix 1.14 |  |  |
| Cell dimensions |  |  |  |
| a=b=c (Å) | 525.7224 | 517.104 | 525.7224 |
| α=β=γ (°) | 90 |  |  |
| Model composition |  |  |  |
| Protein residues | 7,050 | 6,521 | 7,050 |
| Side chains assigned | 7,050 | 6,521 | 7,050 |
| Biotin | 6 | 5 | 6 |
| Acetyl-CoA | 0 | 6 | 0 |
| Propionyl-CoA | 0 | 0 | 6 |
| R.m.s deviations |  |  |  |
| Bonds length (Å) | 0.010 | 0.010 | 0.010 |
| Bonds Angle (°) | 1.064 | 1.008 | 1.041 |
| Ramachandran plot statistics (%) |  |  |  |
| Preferred | 96.58 | 96.55 | 96.43 |
| Allowed | 2.82 | 2.72 | 2.92 |
| Outlier | 0.60 | 0.72 | 0.65 |

**Table S2.** Cryo-EM data collection, reconstruction, refinement, and validation statistics of human MCC structures.

| Sample | MCC-**apo** | MCC-**ACO** | MCC-**PCO** |
| --- | --- | --- | --- |
| **PDB code** | 8XL6 | 8XL7 | 8XL8 |
| **EMDB code** | EMD-38439 | EMD-38440 | EMD-38441 |
| **Data collection** |  |  |  |
| EM equipment | Titan Krios (Thermo Fisher Scientific) |  |  |
| Voltage (kV) | 300 |  |  |
| Detector | Gatan K3 Summit |  |  |
| Energy filter | Gatan GIF Quantum, 20 eV slit |  |  |
| Pixel size (Å) | 1.0773 |  |  |
| Electron dose (e<sup>-</sup>/Å<sup>2</sup>) | 50 |  |  |
| Defocus range (μm) | −1.2 to −2.2 |  |  |
| Number of collected micrographs | 11,481 | 4,036 | 10,898 |
| **3D Reconstruction** |  |  |  |
| Software | cryoSPARC 3.3.2 |  |  |
| Number of used particles | 407,407 | 117,900 | 476,733 |
| Resolution (Å) | 2.29 | 2.85 | 2.36 |
| Symmetry | C1 |  |  |
| Map sharpening B-factor (Å<sup>2</sup>) | 69.9 | 88.4 | 75.7 |
| **Refinement** |  |  |  |
| Software | Phenix 1.14 |  |  |
| **Cell dimensions** |  |  |  |
| a=b=c (Å) | 525.7224 | 517.104 | 525.7224 |
| α=β=γ (°) | 90 |  |  |
| **Model composition** |  |  |  |
| Protein residues | 5,780 | 5,704 | 5,628 |
| Side chains assigned | 5,780 | 5,704 | 5,628 |
| Biotin | 6 | 6 | 6 |
| Acetyl-CoA | 0 | 6 | 0 |
| Propionyl-CoA | 0 | 0 | 6 |
| **R.m.s deviations** |  |  |  |
| Bonds length (Å) | 0.011 | 0.013 | 0.011 |
| Bonds Angle (°) | 1.015 | 1.146 | 1.018 |
| **Ramachandran plot statistics (%)** |  |  |  |
| Preferred | 95.05 | 93.77 | 94.63 |
| Allowed | 4.74 | 5.60 | 4.94 |
| Outlier | 0.21 | 0.64 | 0.43 |

